# Community diversity favors coexistence between bacteria and parasitic bacteriophages

**DOI:** 10.64898/2026.07.21.739906

**Authors:** Michael Blazanin, William An, Abraham B. Tolkoff, Paul E. Turner

## Abstract

Microbial communities often contain diverse bacteria and their specific viruses (bacteriophages). Lytic phages frequently select for host bacteria to evolve resistance, in-turn threatening phage extinction. However, in nature phage-bacteria interactions do not occur in isolation, but within complex communities containing other species. To test how community context can alter resistance evolution and phage persistence, we experimentally evolved *Pseudomonas aeruginosa* bacteria and a *Pseudomonas*-targeting phage with and without a community representing a cystic fibrosis disease pulmonary infection. Phages selected for rapid evolution of receptor-mediated resistance, regardless of community context. However, susceptible bacteria often persisted due to resistance-growth trade-offs. A diverse community favored increased frequencies of susceptibility, fueling higher phage densities and fewer phage extinctions. Our findings show that community complexity can constrain the evolution of phage resistance in bacteria, promoting host-parasite coexistence and fostering microbial diversity.

## Introduction

Microbial communities are a dominant constituent of our biosphere (*1*), with impacts from natural ecosystem function (*2*) to human health (*3*). Within these communities, a complex web of interspecies interactions mediate competition, facilitation, and parasitism, ultimately driving coexistence or extinction. The most numerous members of microbial communities are bacteria and their specific viruses, bacteriophages (*4*). Phages are ubiquitous and can vary greatly in their life history, from lysogens that can be transmitted vertically across generations to obligately lytic parasites that kill their host (*5*, *6*). Indeed, lytic phages can be so effective that they are increasingly important candidates for the treatment of otherwise-untreatable bacterial infections (*7*). Lytic phages pose a severe threat of infection to their bacterial hosts, frequently driving bacteria to evolve resistance against phage attack (*8*). In turn, phages can face strong selection to overcome resistance to maintain infectivity. However, evolutionary interactions between bacteria and phages and the possibilities for these microbes to coexist do not happen in isolation: rather, they are likely to be shaped by the presence of other species in the surrounding community (*9*, *10*). Despite this importance, there have been few direct tests of how community context can alter the evolution and persistence of bacteria and their phages.

The evolution of bacterial resistance to phage infection is multifaceted, with consequences for the survival of both bacterial and phage populations. Bacterial resistance can arise through a variety of mechanisms, one of the most common being mutations to structures on the bacterial cell-surface used by phages as receptors (*11*, *12*). However, resistance often carries fitness costs (i.e. tradeoffs), with examples including reduced growth (*13*, *14*), competitive ability (*15*), or ability to infect an animal host (*16*). Because of these costs, bacterial resistance does not always fix in a population. Instead, phage-resistant and phage-susceptible bacteria may be maintained in a competitive equilibrium (*15*, *17–19*). Alternatively, resistance may spur the evolution of the phage, sparking coevolution by arms race or fluctuating selection dynamics (*20*). Ultimately, while the evolution of bacterial resistance can lead to phage extinction, maintenance of susceptible bacteria or phage counter-adaptation can both support long-term persistence of phage populations (*21*). Both mechanisms of phage persistence, costs of resistance and coevolution, could be impacted by the surrounding biotic community. Despite this, few experiments have directly tested how community context can alter the persistence of phages through shaping the evolution of resistance (*10*, *16*, *22–35*).

To address this gap, here we used experimental evolution with a medium-complexity synthetic community to test how community context alters the evolution and coexistence of a phage and its bacterial host. Such experiments are an emerging approach to reveal the eco-evolutionary dynamics of phage-bacterial communities (*10*, *16*, *22–35*) and bridge the longstanding gap between bottom-up (reductionist) and top-down (holistic) understanding (*36*). In our community we used *Pseudomonas aeruginosa*, a leading cause of opportunistic infections (*37*), and its lytic phage TIVP-H6 (hereafter, H6), a developing therapeutic candidate (*38*, *39*). To emulate the species community context of a pulmonary infection in vulnerable individuals, we used three bacterial species that commonly co-occur in the lungs of people with cystic fibrosis disease: *Staphylococcus aureus*, *Enterococcus faecalis*, and *Achromobacter xylosoxidans* (*40*). We experimentally evolved mixtures of *P. aeruginosa* bacteria and phage, finding that *Pseudomonas* rapidly and universally evolved costly resistance via mutations in genes underlying the phage receptor: the type IV pilus. However, the presence of other bacterial species shifted the balance of susceptible and resistant *Pseudomonas* bacteria, favoring increased susceptibility and enabling higher densities and persistence of the phage in the more diverse community context.

## Materials and Methods

### Strains and culture conditions

We used *Pseudomonas aeruginosa* strain PA14, alongside *Staphylococcus aureus* subspecies Aureus (ATCC #25904), *Enterococcus faecalis* (ATCC #700802), and an *Achromobacter xylosoxidans* strain isolated from a cystic fibrosis patient’s sputum sample. We used phage TIVP-H6 (hereafter, H6) (*38*, *39*), which putatively binds the *P. aeruginosa* type IV pilus, and does not infect the other three bacterial ancestral strains. Unless otherwise noted, bacteria were cultured at 37°C in Bolton Broth Base (GranuCult, EMD Millipore Corporation #1.00068).

### Experimental evolution

Overnight bacterial cultures were grown from a single colony in 10 mL of Bolton Broth Base shaking at 37°C. Then, bacteria and phages were inoculated into 25 mL glass flasks containing 10 mL of Bolton Broth Base. For bacteria, 100 µL of each overnight bacterial culture was added, with replicates sharing the same population number across the four treatments being inoculated from a single flask for each bacterial strain. For phage H6, a phage stock was added to the flask to achieve a final concentration of 10^5^ pfu/mL. All populations were inoculated from the same phage stock. Communities were then incubated statically (non-shaking) at 37°C for 24 hours. After 24 hours, communities were passaged by vortexing thoroughly and sampling 100 µL, which was added to a new flask containing 10 mL Bolton Broth Base. This was repeated for 9 total days of incubation.

After each day of incubation, communities were sampled to make frozen stocks and to estimate population densities. Samples from each community on each day were mixed with glycerol to a final concentration of 25% glycerol and then stored at -80°C. Communities were also sampled and serially diluted in Bolton Broth Base. For each community, 2.5 µL was spotted onto five different agar plates to quantify the density of each member of the community: Pseudomonas isolation agar (DOT Scientific Inc NCM0150A) for *P. aeruginosa* densities, Tryptic Soy Agar (Criterion C7141 Tryptic Soy Broth with 15 g/L agar) with 75 g/L NaCl for *S. aureus* densities, Tryptic Soy Agar with 1 µg/mL ciprofloxacin for *A. xylosoxidans* densities, and Tryptic Soy Agar with 50 µg/mL colistin and 20 µg/mL vancomycin to estimate *E. faecalis* densities. For phage densities, 10 µL chloroform was added to a 100 µL sample and vortexed thoroughly before diluting. Samples were then spotted onto LB (10g Tryptone, 5g Yeast Extract, 10g NaCl, MPBio #3002022) plates with a 4 mL layer of 0.5% agar LB containing 200 µL of an overnight culture of ancestral PA14. Note that spotting revealed that both Population 9 treatments with no phage added had been inadvertently contaminated with phages, and were excluded from analyses.

### Resistance assays

To carry out resistance assays, we first isolated evolved *Pseudomonas* and phages from passages 0 (ancestral population), 1, 2, 4, and 8 days by double-isolation streaking on Pseudomonas agar or double-plaque purification on ancestral *Pseudomonas* lawns. We carried out an initial resistance screen of all 5116 within-population combinations using either a spot test or a cross-streak for each combination. For spot tests, each undiluted phage stock (minimum titer 7.3 × 10^8^ pfu/mL) was spotted onto each bacterial lawn, producing a binary outcome (clearing or no clearing). For cross streaks, 15 µL of each phage stock was spread in a single line across a plate, then each bacterial isolate was streaked across it once.

After this initial screen of all combinations, any phage-bacteria combinations which showed evidence of susceptibility were re-tested with an efficiency of plaquing (EOP) assay: phage stocks were serially diluted and spotted onto Bolton Broth plates with a 4 mL top-layer of 0.5% agar Bolton Broth Base containing 200 µL of an overnight culture of the bacterial isolate of interest. For visualization and analysis purposes, we categorized evolved clones into resistance categories based on an EOP threshold of 0.5: bacterial clones with an average EOP (across all tested phages) higher than 0.5 were labeled susceptible, clones with an EOP lower than 0.5 but still formed plaques or clearings were labeled partially susceptible/resistant, and clones that did not form plaques or clearings (i.e. EOP was below detection) were labeled fully resistant. These categories gave identical results to categorization based on plaque morphology (Fig S7, where clones where phages formed ancestral-looking clear plaques were fully susceptible, clones where phages formed cloudy or tiny plaques were partially susceptible/resistant, and clones where phages formed no plaques/clearing were fully resistant).

### Twitching motility

To quantify twitching motility (*41*), bacterial isolates were grown with overnight shaking in 10 mL Bolton Broth at 37°C. 1 µL of culture was inoculated underneath the agar of a 1 day-old 100mm petri plate containing 10 mL of Bolton media (10 g/L agar) by stabbing a pipette tip to the bottom of the plate. Plates were incubated for 24 hours, the agar removed, and images of the bacterial growth on the plastic of each plate were scanned at 300 dpi. The average radius of each area of bacterial growth was manually measured with Fiji (*42*). We quantified twitching motility for two isolates from each population and each treatment at timepoints 1, 2, 4, and 8 days. In each batch there were two technical replicates of the *pilA* non-motile control strain, and a single technical replicate of every other strain. Twitching rates were batch-corrected by scaling all values such that the ancestral and *pilA* strains in each batch equal their respective global average values.

### Growth curves

Bacterial isolates were grown with overnight shaking in 10 mL Bolton Broth at 37°C. Each culture was normalized to an optical density at 600 nm (OD600) of 0.05, then 200 uL was added to each well of a 96 well plate. Plates were incubated in a Biotek Epoch 2 plate reader at 37°C with shaking and the OD600 read every 15 minutes. Data was analyzed using gcplyr (*43*): blanks were subtracted, then data was smoothed by a moving median of 3 data points followed by a moving average of 5 data points. The cellular growth rate was calculated on log-transformed data with a moving window of 5 data points. Then the lag time, maximum growth rate, maximum density (within 23 hours), and area under the curve (from 0.25 to 23 hours) were calculated. Batch correction was carried out by fitting a mixed effects model of each trait with fixed effects for treatment and timepoint plus random effects for population, isolate, and batch, then subtracting the batch effect.

### Sequencing

To sequence isolated bacterial strains and mixed population/community samples, the -80°C frozen sample or stock (final concentration 25% glycerol) was thawed on ice, then 1 mL removed and spun at maximum speed for 10 minutes to form a cell pellet. 450 uL of Monarch DNA/RNA Protection Reagent (New England Biolabs T2011L) was added (an approximately 1:10 dilution for the cells). Samples were then sent to SeqCoast Genomics (Portsmouth, New Hampshire) for sequencing. SeqCoast lysed samples using MaxMAX Microbiome bead beating tubes, then extracted samples using the DNeasy 96 PowerSoil Pro QIAcube HT Kit.

In order to assemble an ancestral genome to call mutations against, an ancestral isolate of *P. aeruginosa* was long-read sequenced. SeqCoast followed their standard workflow: “Samples were prepared for sequencing using the Oxford Nanopore Technologies SQK-NBD114 native barcoding kit. Long Fragment Buffer was used to promote longer read lengths. Sequencing was performed on either the GridION platform using a FLO-MIN114 Spot-ON Flow Cell (R10 version) or the PromethION 2 Solo platform using a FLO-PRO114M Flow Cell (R10 version). Sequencing was performed with a translocation speed of 400bps. Base calling was performed on the GridION using the ‘super-accurate’ basecalling model with barcode trimming enabled (MinKNOW version: 24.02.16, Dorado version: 7.3.11).”

For all other samples, short-read sequencing was carried out. SeqCoast followed their standard workflow: “Samples were prepared using the Illumina DNA Prep tagmentation kit and Illumina Unique Dual Indexes. Sequencing was performed on the Illumina NextSeq2000 platform using a 300 cycle flow cell kit to produce 2×150bp paired reads. 1-2% PhiX control was spiked into the run to support optimal base calling. Read demultiplexing, read trimming, and run analytics were performed using DRAGEN v4.2.7, an on-board analysis software on the NextSeq2000.”

We carried out sequencing of 1 ancestral *P. aeruginosa* bacterial isolate, the ancestral *A. xylosoxidans* strain, 83 evolved *P. aeruginosa* bacterial isolates (Table S1), and 88 metagenomes of mixed populations/communities: the ancestral *P. aeruginosa* population for each replicate, the entire community at timepoint day 8 for all treatments, and the entire community at timepoints 1, 2, and 4 days for phage-added treatments.

The ancestral *A. xylosoxidans* genome was assembled from short reads using shovill v1.1.0 (*44*). The ancestral *P. aeruginosa* genome was assembled from long reads using Autocycler v0.5.2 (*45*). Both ancestral genomes were annotated using bakta v1.11.4 (database v6.0) (*46*). Completeness of the ancestral genomes was confirmed using checkm2 v1.1.0 (*47*). The ancestral phage H6 genome was downloaded from GenBank: PP203295.1. The ancestral *S. aureus* (#25904, version published 2019-08-27) and *E. faecalis* (#700802, version published 2019-05-14) genomes were downloaded from the ATCC Genome Portal Database.

Sample sequences were filtered using fastp v1.0.1 on default settings (*48–50*). Mutations in samples were called using breseq v0.39.0 (*51–53*): isolates were called using default settings in consensus mode; populations/communities were called in polymorphism mode with maximum-read-mismatches set to 3, but otherwise default settings. *P. aeruginosa* isolates and timepoint 0 *P. aeruginosa* populations were called against the PA14 reference, while all other samples were called against all 5 reference genomes.

Genes related to pilus function were identified manually from gene names, annotations, and GO terms (see Supplemental Materials). To calculate the total frequency of all mutations in pilus-related genes without linkage data, we assume that distinct pilus mutations arise on separate genetic backgrounds and are competing with one another (see Figs S17, S18 for an alternate assumption), consistent with our isolate sequencing where very few isolates have multiple pilus mutations (Fig S8).

## Results

To test how community diversity alters the coevolution and coexistence of phages and their bacterial hosts, we carried out experimental evolution of *P. aeruginosa* in the presence or absence of phage H6 and the presence or absence of three additional bacterial species. We found that community context strongly altered the densities of phage H6, while populations of all four bacterial species converged on similar equilibrium densities regardless of treatment (Fig 1). In the low-diversity communities, phage populations were significantly less dense (linear model of phage density days 2 – 9 with each day and treatment as factors, effect of treatment t = 3.924, p < 0.001), with half of the populations dropping below 10^4^ pfu/mL at some point in the experiment. In contrast, in the high-diversity communities, most phage population densities remained high, with only one population ever dropping below 10^4^ pfu/mL.

**Figure 1.**
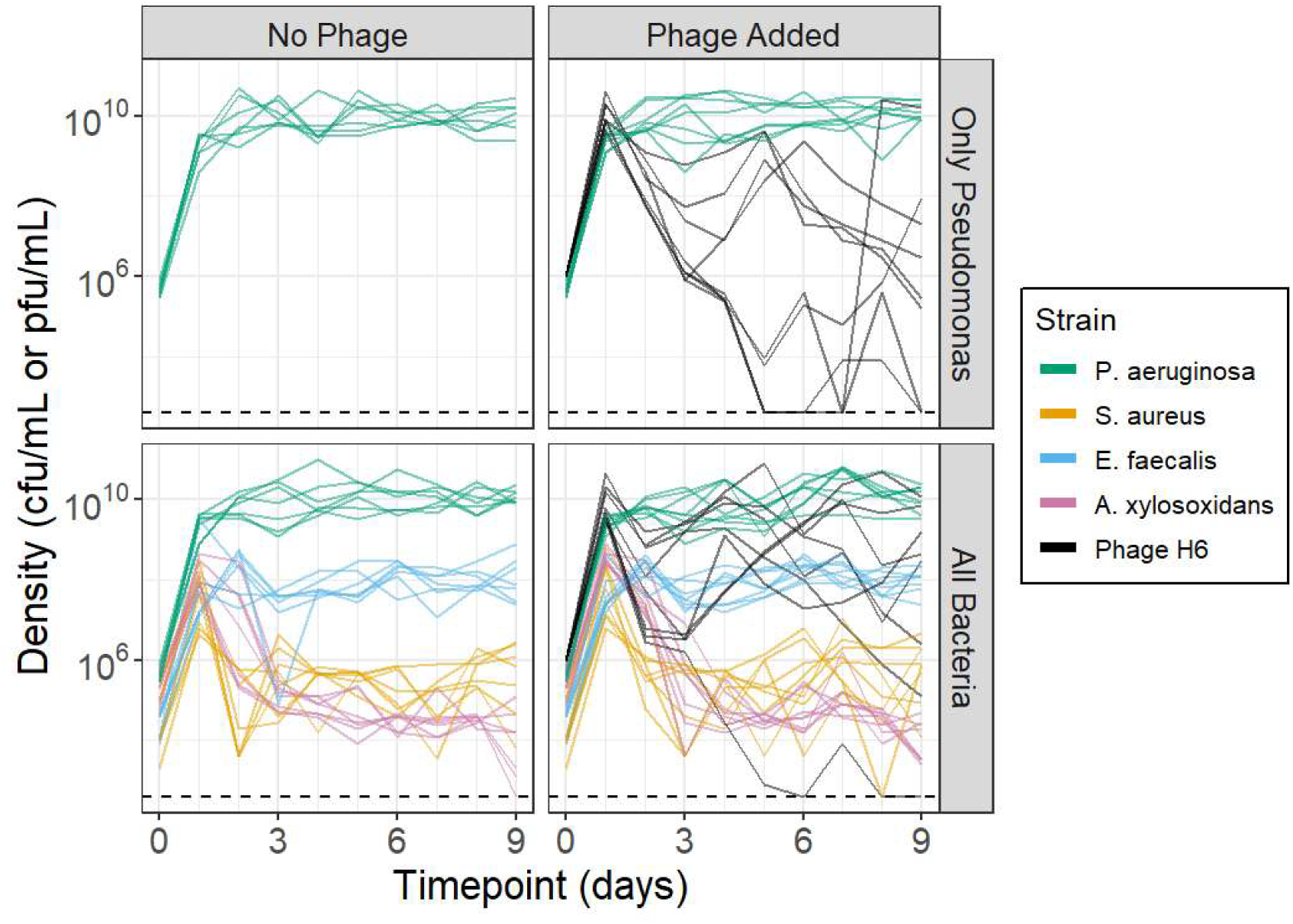
Population dynamics during experimental evolution show that bacterial diversity favors higher phage densities and more frequent phage persistence. Mixed communities of bacteria and phages were passaged daily for 9 days with a 1:100 dilution, with population densities measured by plating. Each line is an independently evolving replicate population.

To test whether differences in phage density could be explained by differences in the evolution of bacterial resistance or phage infectivity, we isolated evolved strains of *P. aeruginosa* and phage H6 from each population at timepoints 0, 1, 2, 4, and 8 days. We quantified the resistance of all 5116 combinations of phage and bacteria within each population. We observed no evidence of phage evolution in any population, including no evidence of any evolved host switch to infect the other bacterial species. In contrast, we observed rapid evolution of bacterial resistance among *Pseudomonas* isolates in all populations (Fig 2). Despite widespread resistance, partially and even fully phage-susceptible bacteria did persist in many populations. However, there was no detectable difference between treatments in the frequency of susceptibility among bacterial isolates (linear model of resistance frequency within each population by community treatment and timepoint, ANOVA effect of treatment intercept p = 0.18, effect of treatment slope with time p = 0.59).

**Figure 2.**
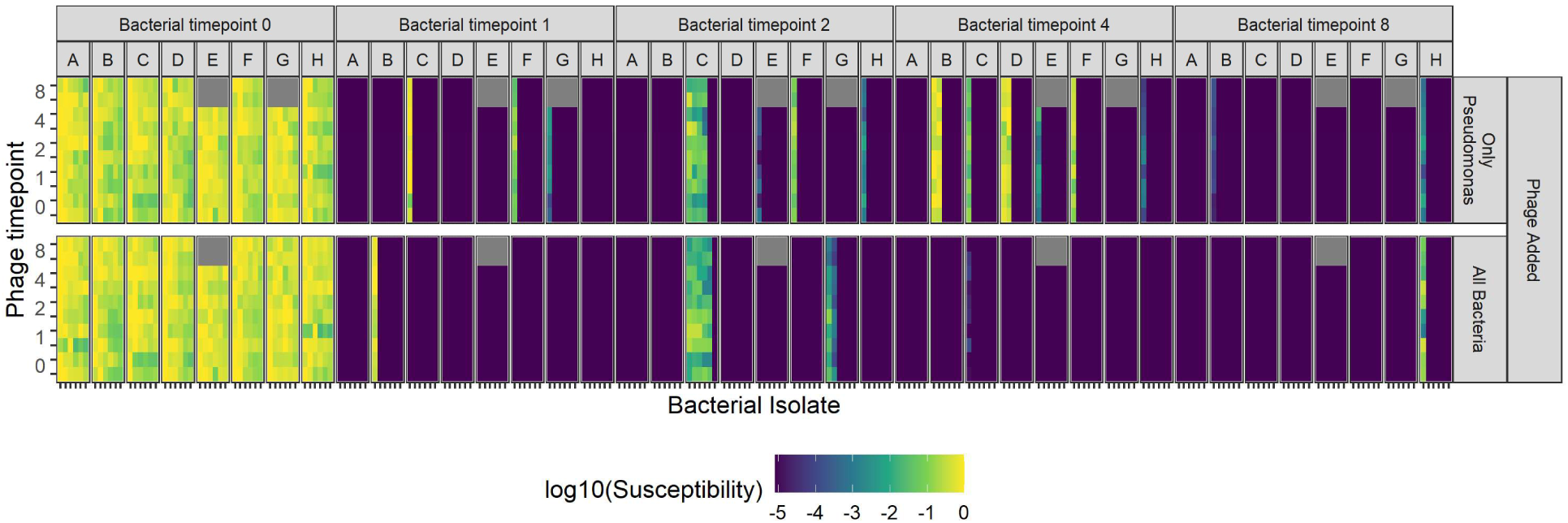
Infectivity/Resistance dynamics of phage H6 and *P. aeruginosa* reveal widespread resistance evolution regardless of community context. Six bacterial and two phage clones were isolated from each population (A through H) at timepoints 0, 1, 2, 4, and 8 days. The susceptibility/resistance of all 5116 combinations was quantified, with timepoints where no phage could be isolated due to phage extinction plotted in gray. Susceptibility is the efficiency of plaquing, relative to the bacterial strain which showed the highest number of plaques for each phage strain in each batch.

To identify the genetic bases for phage resistance in our experiment, we sequenced at least one evolved bacterial isolate from each phage-added population at timepoints 1, 2, 4 and 8 days (83 isolates total). H6 attacks *P. aeruginosa* using the type IV pilus as a receptor (*54*), and indeed nearly all phage-resistant bacteria had one or more mutations in pilus genes, while none of the fully-susceptible bacteria had mutations in any pilus genes (Fig S8). However, there were no substantial differences between treatments. We next tested how widespread pilus mutations are, and whether pili functionality correlated with phage susceptibility. Mutations that modify or break the type IV pilus might confer resistance but also compromise pilus function, so we inferred pilus function by quantifying pilus-dependent twitching motility. Among a larger collection of two isolates from each population and timepoint, we observed rapid evolution of lost motility in the presence of phages, to levels equal to a pilus knockout (Δ*pilA*) control [Fig 3A, ANOVA for inclusion of treatment factor on fit of mixed effect model for batch-corrected twitching, with fixed effects for each treatment (intercept) and interaction between treatment and numeric timepoint (slope), and random effect for population, p < 0.001]. Evolved clones showed significant differences in twitching motility depending on their level of phage-susceptibility (Fig 3B, ANOVA for inclusion of resistance category factor on fit of mixed effect model for batch-corrected twitching, with fixed effects for phage treatment, community treatment, and resistance category and random effect for population, p < 0.001). However, there was no difference in twitching motility between *Pseudomonas*-only and all-bacteria treatments (Fig 3B, ANOVA for inclusion of community treatment factor on fit of previous model, p = 0.69). Thus, in our experiment *P. aeruginosa* evolved phage resistance via mutations to the phage receptor.

**Figure 3.**
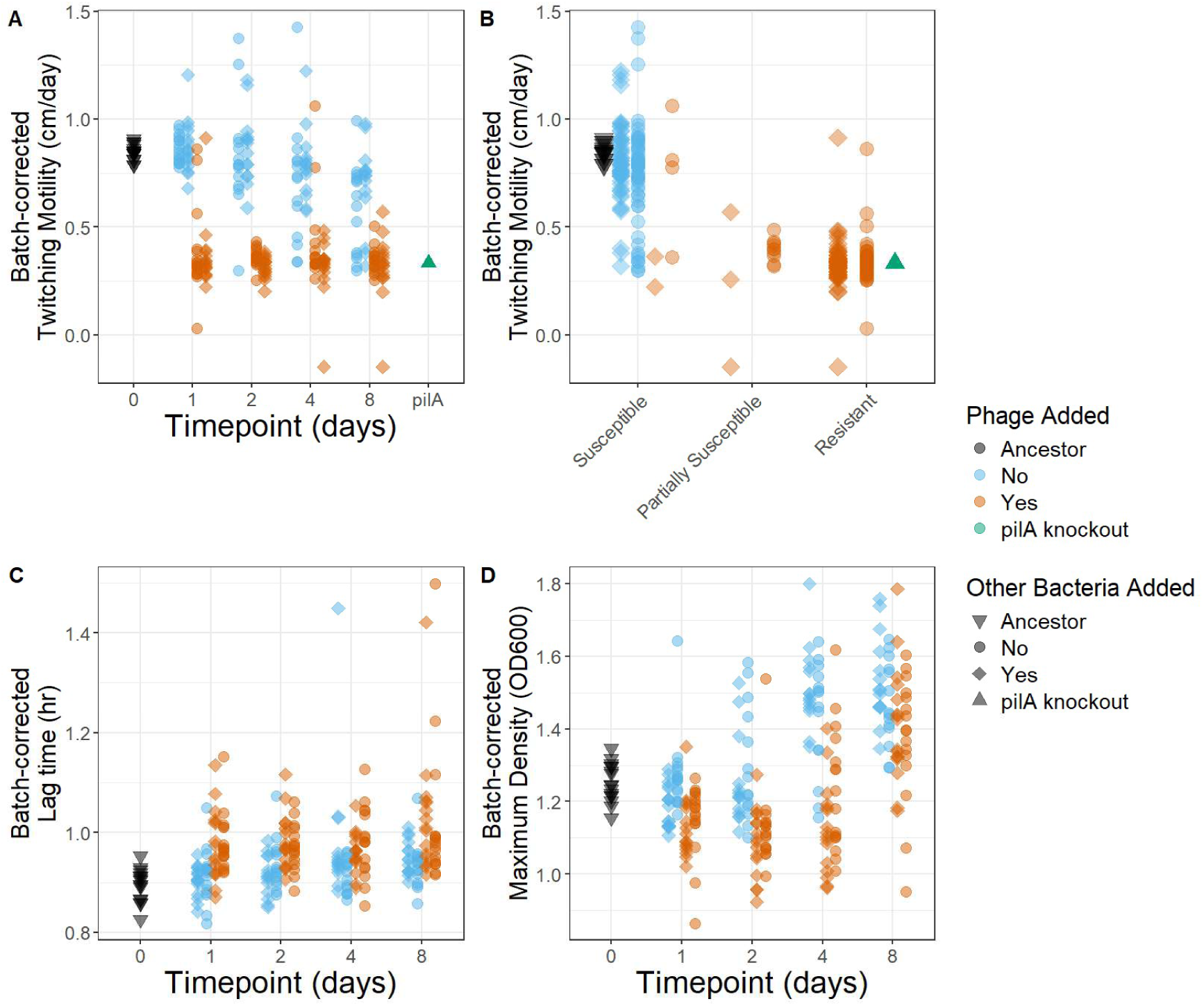
Phage-exposed *P. aeruginosa* rapidly evolve loss of pilus function and show phenotypic costs of resistance, regardless of community treatment. **A.** Twitching motility was quantified for two evolved clones from each population and timepoint. Batch-correction was carried out relative to the ancestral and pilA isolates, which were run in every batch. **B.** Resistance was categorized by quantitative efficiency of plaquing and phage plaque morphology. **C, D.** For each of two evolved clones from each population and timepoint, we used growth curves to quantify lag time and maximum density.

We next tested whether evolved bacterial clones paid any costs (relative to the ancestor) or opportunity costs (relative to bacteria evolving in the absence of phages or other bacteria). For the same collection of two isolates from each population and timepoint, we used growth curves to quantify lag time and maximum density. Generally, bacteria from phage-added treatments had longer lag times and lower maximum densities than bacteria from no-phage treatments (Fig 3C, 3D, S11), suggesting reduced growth traits in these bacteria. However, the dynamics of these traits over time differed between traits. In both traits bacteria show an immediate cost of evolution in timepoint 1, relative to the ancestor. However, in subsequent timepoints no-phage bacteria began to evolve increased maximum density, with phage-added bacteria following the same trend but paying an ‘opportunity cost’ and lagging behind. Motivated by previously-published findings (*55–58*), we also quantified several other traits, but generally found no differences between treatments, including biofilm formation and pyocyanin and pyoverdine production (Figs S12, S13). Generally, these data reflect both costs and opportunity costs due to rapid evolution of phage resistance in our experiment.

Next, we tested how phage and community treatments altered the molecular evolution of the *P. aeruginosa* populations. Multiple mutations were present in each population, with an enrichment of pilus-related mutations in phage-added treatments (Fig 4A). As expected, pilus mutations were significantly more frequent in phage-added treatments (Fig S17, ANOVA for inclusion of phage treatment factor on fit of mixed effect model for total frequency at timepoint day 8, with fixed-effect for each treatment and random effect for population, p < 0.001). In several populations we also observed turnover in the dominant pilus mutation over time (Fig 4B), suggesting competition between lineages with different pilus mutations.

**Figure 4.**
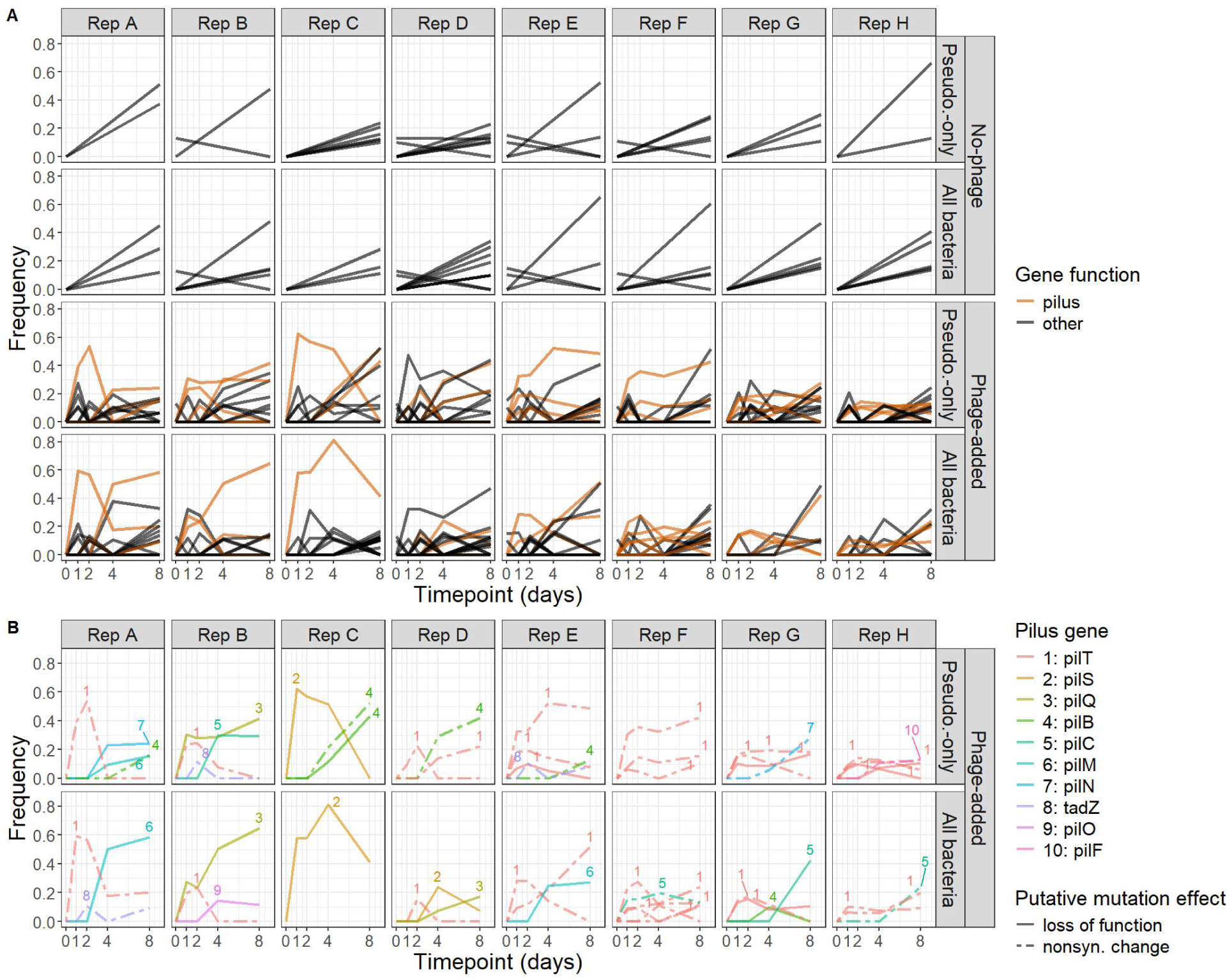
Molecular evolution in populations in phage-added treatment is dominated by mutations in diverse pilus-related genes. Whole communities in no-phage treatments were sequenced at timepoints 0 and 8, while communities in phage-added treatments were sequenced at timepoints 0, 1, 2, 4, and 8. Each line is a unique mutation observed over time. For visualization only mutations which exceed 10% frequency at some point are plotted here. **A.** Mutations are colored by whether they affect a pilus-related gene. **B.** Mutations are colored and numbered for convenience by gene. Mutations are categorized as putatively loss of function (frameshift indels and nonsense SNPs) or nonsynonymous change (in-frame indels and nonsynonymous SNPs).

We tested whether the molecular evolutionary dynamics of *P. aeruginosa* could explain the divergent population dynamics of phages observed during experimental evolution (Fig 1). We found that phage density was strongly and significantly negatively correlated with total pilus mutation frequency (Fig 5A, linear model of log_10_ phage density by treatment × pilus mutation frequency, only-Pseudomonas coefficient = -3.3, p = 0.04; all-bacteria coefficient = -3.1, p = 0.05, but see Fig S18 for an alternate assumption). Moreover, pilus mutation frequencies were moderately but significantly higher in the Pseudomonas-only treatment (Fig 5B, Pseudomonas-only effect of 0.12 with p = 0.027). We also carried out a GO term enrichment analysis, which produced consistent results with this effect: pilus-related terms were enriched in mutations in the Pseudomonas-only treatments relative to mutations in the all-bacteria treatments (Table S2). These results suggest that: 1) phage density is being supported by the size of the population of bacteria lacking pilus mutations (Fig 5A), and 2) the addition of other bacterial species favored higher frequencies of *Pseudomonas* bacteria with wild type pili (Fig 5B).

**Figure 5.**
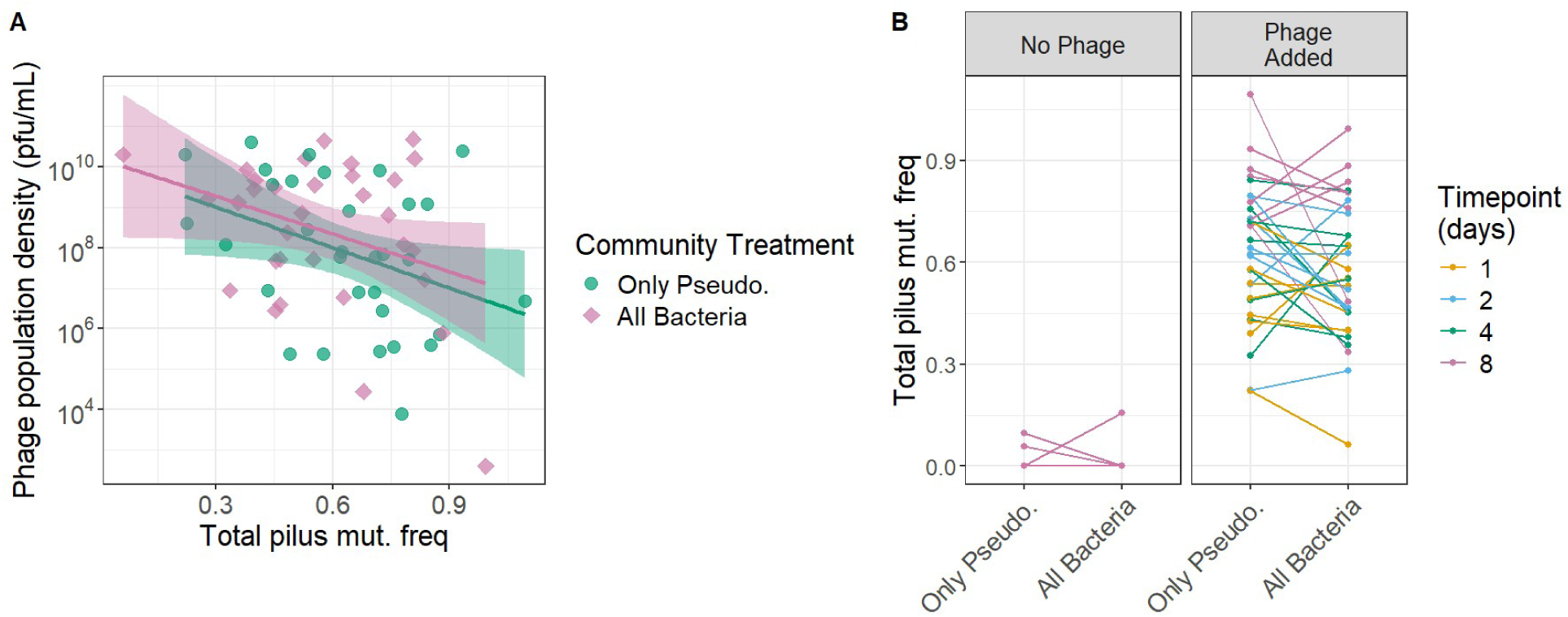
Mutations in pilus-related genes predict lower phage density and are more frequent in the Pseudomonas-only treatment. The total frequency of all mutations in pilus-related genes was calculated assuming that mutations arise on separate genetic backgrounds and are competing with one another (see Figs S17, S18 for an alternate assumption). **A.** Each point is a replicate population at a single timepoint. **B.** Lines link frequencies from matched replicate populations across treatments.

## Discussion

Here we tested how microbial community diversity can alter the evolution of bacterial resistance to phages and the persistence of bacteria and phages. Our results are the first to show that community diversity can interact with trade-offs to favor phage-bacteria coexistence. In our experiments we found that phages drove the rapid evolution of widespread bacterial resistance to phage infection (Fig 2) via mutations in genes underlying the phage binding receptor, the type-IV pilus (Figs 3A, 3B, 4, S8). However, susceptible bacteria were able to persist (Figs 2, 5A), likely because resistance was costly, reflecting a trade-off between growth and resistance to phage infection (Figs 3C, 3D, S11). Species diversity in the community interacted with this trade-off, favoring higher frequencies of phage-susceptible bacteria (Fig 5B), supporting higher phage densities and fewer phage extinctions (Figs 1, 5A).

Our work suggests that divergent past findings on the effects of community diversity on phages and bacteria might be unified by characterizing how community context interacts with phage resistance trade-offs. For example, previous work has diverged on whether community diversity constrains (*16*, *22*, *24*, *34*) or has no effect on (*23*, *25–27*, *30*, *32*, *33*) the bacterial evolution of phage resistance. Existing work has also diverged on whether community diversity increases (*32*, *34*), decreases (*24*, *34*), or has no effect on (*29*) phage-bacteria coexistence. These discrepancies might be explained by differences in the trade-offs associated with phage resistance in each case. Indeed, both experimental and theoretical work has shown that trade-offs associated with the evolution of phage resistance can underlie coexistence between phages and bacteria in simple communities (*15*, *17–19*, *59*, *60*). However, until now the role of phage resistance trade-offs in more complex microbial communities had remained largely untested. Our findings suggest that the fitness costs of phage resistance and the coexistence of phage and bacteria may both be heightened by the species diversity in natural microbial communities.

Our work is also the first to experimentally test how multispecies bacterial communities might alter the efficacy of therapeutic phage steering in vitro. Phage steering is the use of phage therapy to treat bacterial infections where the evolution of resistance has beneficial trade-offs, for example the loss of virulence or antibiotic resistance (*7*, *61*). While phage steering is a frequent aim of phage therapy (*62*), including with H6 (*38*, *39*), demonstrations of phage steering come overwhelmingly from in vitro experiments with a single bacterial species (*61*) or in vivo tests where mechanisms can be difficult to untangle (*39*). The only exception we are aware of found that phage steering of *P. aeruginosa* was not compromised by the presence of *S. aureus* (*30*). Our experiments are the first to test phage steering in an in vitro community of more than two species, and our results strengthen the case for phage steering and phage therapy when applied to multispecies infections. In our experiments, *P. aeruginosa* evolved mutations in the phage receptor regardless of community diversity. Moreover, these mutations compromised type IV pilus function (Fig 3A, 3B), changes that would be associated with the loss of virulence in a mammalian host (*39*, *63*). Additionally, multispecies diversity favored persistence of phage populations for longer periods of time, suggesting that phages delivered therapeutically might continue to suppress or steer bacterial pathogens for longer when applied in a diverse community.

Our results may represent a conservative estimate of the possible effects of community context. In many past experiments, the presence of additional bacterial diversity suppressed the density of bacteria and phages (*10*, *16*, *25–27*, *29*, *31*, *33*, *34*). In contrast, in our experiments community diversity had no substantial effect on the density of *Pseudomonas* bacteria, with *Pseudomonas* competitively dominating the community from days 3 to 9. This *Pseudomonas* dominance reflects the cystic fibrosis disease pulmonary infections we were modeling (*64*), and is the context where phage H6 is currently being used therapeutically (*38*, *39*). Despite the dominance of *Pseudomonas*, community context still had strong effects (Figs 1, 5). One possible explanation is that even rare individuals of non-*Pseudomonas* species were sufficient to affect the evolution of resistance (*65*). Alternatively, community context may have altered the trajectory of evolution early in the experiment before *Pseudomonas* reached dominance, consistent with recent work showing that the initial conditions where phage resistance evolves can have lasting effects on long-term resistance strategies (*66*). Regardless of the explanation, our results suggest that community context could be shaping the evolution of phage resistance even when a single bacterial species dominates the community by an order of magnitude or more.

Broadly, our understanding of microbial ecology and evolution has been built from reductionist experiments in monoculture or holistic studies of highly complex communities (*36*). These approaches provide complementary insights, but leave a gap between mechanistic understanding and natural context. This has been especially true in our understanding of phage-bacteria ecology and evolution (*10*). Here, we’ve used experiments with a bacterial community to show that diversity can constrain evolution, promoting the coexistence of phages and their bacterial hosts. Ultimately, these patterns suggest that microbial community diversity begets diversity by maintaining opportunities for ongoing competition and parasitism.

## Acknowledgements

We thank Alita Burmeister, Caroline Turner, David Vasseur, Alvaro Sanchez, Andrew Goodman, and C. Brandon Ogbunugafor for feedback on drafts of this manuscript. We thank Dallas Mould and Jyot Antani for assistance with the twitching motility assay, and Ian Will for assistance with the GO terms analysis. We thank the Yale STARS program for funding supporting WA during work on this project, and the Howard Hughes Medical Institute for funding supporting MB during work on this project.

## References

1. W. B. Whitman, D. C. Coleman, W. J. Wiebe, Prokaryotes: The unseen majority. Proceedings of the National Academy of Sciences 95, 6578–6583 (1998).

2. P. G. Falkowski, T. Fenchel, E. F. Delong, The Microbial Engines That Drive Earth’s Biogeochemical Cycles. Science 320, 1034–1039 (2008).

3. C. Huttenhower, D. Gevers, R. Knight, S. Abubucker, J. H. Badger, A. T. Chinwalla, H. H. Creasy, A. M. Earl, M. G. FitzGerald, R. S. Fulton, M. G. Giglio, K. Hallsworth-Pepin, E. A. Lobos, R. Madupu, V. Magrini, J. C. Martin, M. Mitreva, D. M. Muzny, E. J. Sodergren, J. Versalovic, A. M. Wollam, K. C. Worley, J. R. Wortman, S. K. Young, Q. Zeng, K. M. Aagaard, O. O. Abolude, E. Allen-Vercoe, E. J. Alm, L. Alvarado, G. L. Andersen, S. Anderson, E. Appelbaum, H. M. Arachchi, G. Armitage, C. A. Arze, T. Ayvaz, C. C. Baker, L. Begg, T. Belachew, V. Bhonagiri, M. Bihan, M. J. Blaser, T. Bloom, V. Bonazzi, J. Paul Brooks, G. A. Buck, C. J. Buhay, D. A. Busam, J. L. Campbell, S. R. Canon, B. L. Cantarel, P. S. G. Chain, I.-M. A. Chen, L. Chen, S. Chhibba, K. Chu, D. M. Ciulla, J. C. Clemente, S. W. Clifton, S. Conlan, J. Crabtree, M. A. Cutting, N. J. Davidovics, C. C. Davis, T. Z. DeSantis, C. Deal, K. D. Delehaunty, F. E. Dewhirst, E. Deych, Y. Ding, D. J. Dooling, S. P. Dugan, W. Michael Dunne, A. Scott Durkin, R. C. Edgar, R. L. Erlich, C. N. Farmer, R. M. Farrell, K. Faust, M. Feldgarden, V. M. Felix, S. Fisher, A. A. Fodor, L. J. Forney, L. Foster, V. Di Francesco, J. Friedman, D. C. Friedrich, C. C. Fronick, L. L. Fulton, H. Gao, N. Garcia, G. Giannoukos, C. Giblin, M. Y. Giovanni, J. M. Goldberg, J. Goll, A. Gonzalez, A. Griggs, S. Gujja, S. Kinder Haake, B. J. Haas, H. A. Hamilton, E. L. Harris, T. A. Hepburn, B. Herter, D. E. Hoffmann, M. E. Holder, C. Howarth, K. H. Huang, S. M. Huse, J. Izard, J. K. Jansson, H. Jiang, C. Jordan, V. Joshi, J. A. Katancik, W. A. Keitel, S. T. Kelley, C. Kells, N. B. King, D. Knights, H. H. Kong, O. Koren, S. Koren, K. C. Kota, C. L. Kovar, N. C. Kyrpides, P. S. La Rosa, S. L. Lee, K. P. Lemon, N. Lennon, C. M. Lewis, L. Lewis, R. E. Ley, K. Li, K. Liolios, B. Liu, Y. Liu, C.-C. Lo, C. A. Lozupone, R. Dwayne Lunsford, T. Madden, A. A. Mahurkar, P. J. Mannon, E. R. Mardis, V. M. Markowitz, K. Mavromatis, J. M. McCorrison, D. McDonald, J. McEwen, A. L. McGuire, P. McInnes, T. Mehta, K. A. Mihindukulasuriya, J. R. Miller, P. J. Minx, I. Newsham, C. Nusbaum, M. O’Laughlin, J. Orvis, I. Pagani, K. Palaniappan, S. M. Patel, M. Pearson, J. Peterson, M. Podar, C. Pohl, K. S. Pollard, M. Pop, M. E. Priest, L. M. Proctor, X. Qin, J. Raes, J. Ravel, J. G. Reid, M. Rho, R. Rhodes, K. P. Riehle, M. C. Rivera, B. Rodriguez-Mueller, Y.-H. Rogers, M. C. Ross, C. Russ, R. K. Sanka, P. Sankar, J. Fah Sathirapongsasuti, J. A. Schloss, P. D. Schloss, T. M. Schmidt, M. Scholz, L. Schriml, A. M. Schubert, N. Segata, J. A. Segre, W. D. Shannon, R. R. Sharp, T. J. Sharpton, N. Shenoy, N. U. Sheth, G. A. Simone, I. Singh, C. S. Smillie, J. D. Sobel, D. D. Sommer, P. Spicer, G. G. Sutton, S. M. Sykes, D. G. Tabbaa, M. Thiagarajan, C. M. Tomlinson, M. Torralba, T. J. Treangen, R. M. Truty, T. A. Vishnivetskaya, J. Walker, L. Wang, Z. Wang, D. V. Ward, W. Warren, M. A. Watson, C. Wellington, K. A. Wetterstrand, J. R. White, K. Wilczek-Boney, Y. Wu, K. M. Wylie, T. Wylie, C. Yandava, L. Ye, Y. Ye, S. Yooseph, B. P. Youmans, L. Zhang, Y. Zhou, Y. Zhu, L. Zoloth, J. D. Zucker, B. W. Birren, R. A. Gibbs, S. K. Highlander, B. A. Methé, K. E. Nelson, J. F. Petrosino, G. M. Weinstock, R. K. Wilson, O. White, The Human Microbiome Project Consortium, Structure, function and diversity of the healthy human microbiome. Nature 486, 207–214 (2012).

4. A. R. Mushegian, Are there 10^31 virus particles on Earth, or more, or less? Journal of Bacteriology 202, e00052–20 (2020).

5. Z. Hobbs, S. T. Abedon, Diversity of phage infection types and associated terminology: the problem with “Lytic or lysogenic.” FEMS Microbiology Letters 363, 1–8 (2016).

6. S. Mäntynen, E. Laanto, H. M. Oksanen, M. M. Poranen, S. L. Díaz-Muñoz, Black box of phage–bacterium interactions: exploring alternative phage infection strategies. Open Biology 11, 210188 (2021).

7. K. E. Kortright, B. K. Chan, J. L. Koff, P. E. Turner, Phage Therapy: A Renewed Approach to Combat Antibiotic-Resistant Bacteria. Cell Host and Microbe 25, 219–232 (2019).

8. M. D. Hall, G. Bento, D. Ebert, The Evolutionary Consequences of Stepwise Infection Processes. Trends in Ecology & Evolution 32, 612–623 (2017).

9. A. Betts, C. Rafaluk, K. C. King, Host and Parasite Evolution in a Tangled Bank. Trends in Parasitology 32, 863–873 (2016).

10. M. Blazanin, P. E. Turner, Community context matters for bacteria-phage ecology and evolution. ISME J 15, 3119–3128 (2021).

11. S. J. Labrie, J. E. Samson, S. Moineau, Bacteriophage resistance mechanisms. Nature Reviews Microbiology 8, 317–327 (2010).

12. K. D. Seed, Battling Phages: How Bacteria Defend against Viral Attack. PLOS Pathogens 11, e1004847 (2015).

13. B. J. M. Bohannan, R. E. Lenski, The relative importance of competition and predation varies with productivity in a model community. The American Naturalist 156, 329–340 (2000).

14. B. J. M. Bohannan, B. Kerr, C. M. Jessup, J. B. Hughes, G. Sandvik, Trade-offs and coexistence in microbial microcosms. Antonie Van Leeuwenhoek 81, 107–115 (2002).

15. L. Chao, B. R. Levin, F. M. Stewart, A Complex Community in a Simple Habitat: An Experimental Study with Bacteria and Phage. Ecology 58, 369–378 (1977).

16. E. O. Alseth, E. Pursey, A. M. Lujan, I. McLeod, C. Rollie, E. R. Westra, Bacterial biodiversity drives the evolution of CRISPR-based phage resistance in *Pseudomonas aeruginosa*. Nature 574, 549–574 (2019).

17. B. Koskella, M. A. Brockhurst, Bacteria-phage coevolution as a driver of ecological and evolutionary processes in microbial communities. FEMS Microbiology Reviews 38, 1–16 (2014).

18. B. J. M. Bohannan, R. E. Lenski, Linking genetic change to community evolution: Insights from studies of bacteria and bacteriophage. Ecology Letters 3, 362–377 (2000).

19. B. R. Levin, F. M. Stewart, L. Chao, Resource-Limited Growth, Competition, and Predation: A Model and Experimental Studies with Bacteria and Bacteriophage. The American Naturalist 111, 3–24 (1977).

20. S. Gandon, A. Buckling, E. Decaestecker, T. Day, Host–parasite coevolution and patterns of adaptation across time and space. Journal of Evolutionary Biology 21, 1861–1866 (2008).

21. B. Koskella, Resistance gained, resistance lost: An explanation for host–parasite coexistence. PLOS Biology 16, e3000013 (2018).

22. R. Mumford, V. P. Friman, Bacterial competition and quorum-sensing signalling shape the eco-evolutionary outcomes of model in vitro phage therapy. Evolutionary Applications 10, 161–169 (2017).

23. M. Middelboe, A. Hagström, N. Blackburn, B. Sinn, U. Fischer, N. H. Borch, J. Pinhassi, K. Simu, M. G. Lorenz, Effects of bacteriophages on the population dynamics of four strains of pelagic marine bacteria. Microbial Ecology 42, 395–406 (2001).

24. J. Johnke, M. Baron, M. de Leeuw, A. Kushmaro, E. Jurkevitch, H. Harms, A. Chatzinotas, A generalist protist predator enables coexistence in multitrophic predator-prey systems containing a phage and the bacterial predator Bdellovibrio. Frontiers in Ecology and Evolution 5, 1–12 (2017).

25. P. Gómez, A. Buckling, Bacteria-phage antagonistic coevolution in soil. Science 332, 106– 109 (2011).

26. P. Gómez, A. Buckling, Coevolution with phages does not influence the evolution of bacterial mutation rates in soil. ISME Journal 7, 2242–2244 (2013).

27. P. Gómez, A. Buckling, Real-time microbial adaptive diversification in soil. Ecology Letters 16, 650–655 (2013).

28. L. De Sordi, V. Khanna, L. Debarbieux, The Gut Microbiota Facilitates Drifts in the Genetic Diversity and Infectivity of Bacterial Viruses. Cell Host and Microbe 22, 801–808 (2017).

29. R. Sanchez-Martinez, E. Rubio-Portillo, L. Medina-Ruiz, J. Sarasa, M. Enciso, F. Santos, J. Antón, Biological context modulates virus-host dynamics and diversification. bioRxiv [Preprint] (2026). 10.1101/2025.09.10.675297.

30. S. Czerwinski, J. Gurney, Phage steering in the presence of a competing bacterial pathogen. Microbiology Spectrum 13, e02882–24 (2025).

31. T. C. J. Spencer-Drakes, A. Sarabia, G. Heussler, E. C. Pierce, M. Morin, S. Villareal, R. J. Dutton, Phage resistance mutations affecting the bacterial cell surface increase susceptibility to fungi in a model cheese community. ISME Commun 4, ycae101 (2024).

32. R. Dey, A. R. Coenen, N. E. Solonenko, M. N. Burris, A. I. Mackey, J. Galasso, C. L. Sun, D. Demory, D. Muratore, S. J. Beckett, M. B. Sullivan, J. S. Weitz, Density-dependent feedback and higher-order interactions enable coexistence in phage–bacteria community dynamics. ISME J 20, wrag041 (2026).

33. Z. Erdos, D. Padfield, E. Hesse, A. Buckling, M. Castledine, Community Complexity Does Not Weaken Pairwise Coevolution in a Soil Bacterial Community. Ecology Letters 28, e70260 (2025).

34. M. Castledine, D. Padfield, M. Schoeman, A. Berry, A. Buckling, Bacteria–phage (co)evolution is constrained in a synthetic community across multiple bacteria–phage pairs. Microbiology 171, 001577 (2025).

35. L. De Sordi, M. Lourenço, L. Debarbieux, “I will survive”: A tale of bacteriophage-bacteria coevolution in the gut. Gut Microbes 10, 92–99 (2018).

36. R. Tecon, S. Mitri, D. Ciccarese, D. Or, J. R. van der Meer, D. R. Johnson, Bridging the Holistic-Reductionist Divide in Microbial Ecology. MSystems 4, e00265–18 (2019).

37. US Centers for Disease Control and Prevention, “Antibiotic Resistance Threats in the United States, 2019” (2019).

38. M. K. Kim, Q. Chen, A. Echterhof, N. Pennetzdorfer, R. C. McBride, N. Banaei, E. B. Burgener, C. E. Milla, P. L. Bollyky, A blueprint for broadly effective bacteriophage-antibiotic cocktails against bacterial infections. Nat Commun 15, 9987 (2024).

39. B. K. Chan, G. L. Stanley, K. E. Kortright, A. C. Vill, M. Modak, I. M. Ott, Y. Sun, S. Würstle, C. N. Grun, B. I. Kazmierczak, G. Rajagopalan, Z. M. Harris, C. J. Britto, J. Stewart, J. S. Talwalkar, C. R. Appell, N. Chaudary, S. K. Jagpal, R. Jain, A. Kanu, B. S. Quon, J. M. Reynolds, C. C. Teneback, Q.-A. Mai, V. Shabanova, P. E. Turner, J. L. Koff, Personalized inhaled bacteriophage therapy for treatment of multidrug-resistant Pseudomonas aeruginosa in cystic fibrosis. Nat Med 31, 1494–1501 (2025).

40. S. O’Brien, J. L. Fothergill, The role of multispecies social interactions in shaping Pseudomonas aeruginosa pathogenicity in the cystic fibrosis lung. FEMS Microbiology Letters 364, 1–10 (2017).

41. L. Turnbull, C. B. Whitchurch, “Motility Assay: Twitching Motility” in Pseudomonas Methods and Protocols, A. Filloux, J.-L. Ramos, Eds. (Springer, New York, NY, 2014; 10.1007/978-1-4939-0473-0_9)*Methods in Molecular Biology*, pp. 73–86.

42. J. Schindelin, I. Arganda-Carreras, E. Frise, V. Kaynig, M. Longair, T. Pietzsch, S. Preibisch, C. Rueden, S. Saalfeld, B. Schmid, J.-Y. Tinevez, D. J. White, V. Hartenstein, K. Eliceiri, P. Tomancak, A. Cardona, Fiji: an open-source platform for biological-image analysis. Nat Methods 9, 676–682 (2012).

43. M. Blazanin, gcplyr: an R package for microbial growth curve data analysis. BMC Bioinformatics 25, 232 (2024).

44. Torsten Seemann, Shovill, version 1.1.0 (2016); https://github.com/tseemann/shovill.

45. R. R. Wick, B. P. Howden, T. P. Stinear, Autocycler: long-read consensus assembly for bacterial genomes. Bioinformatics 41, btaf474 (2025).

46. O. Schwengers, L. Jelonek, M. A. Dieckmann, S. Beyvers, J. Blom, A. Goesmann, Bakta: rapid and standardized annotation of bacterial genomes via alignment-free sequence identification. Microbial Genomics 7, 000685 (2021).

47. A. Chklovski, D. H. Parks, B. J. Woodcroft, G. W. Tyson, CheckM2: a rapid, scalable and accurate tool for assessing microbial genome quality using machine learning. Nat Methods 20, 1203–1212 (2023).

48. S. Chen, Y. Zhou, Y. Chen, J. Gu, fastp: an ultra-fast all-in-one FASTQ preprocessor. Bioinformatics 34, i884–i890 (2018).

49. S. Chen, fastp 1.0: An ultra-fast all-round tool for FASTQ data quality control and preprocessing. iMeta 4, e70078 (2025).

50. S. Chen, Ultrafast one-pass FASTQ data preprocessing, quality control, and deduplication using fastp. iMeta 2, e107 (2023).

51. J. E. Barrick, G. Colburn, D. E. Deatherage, C. C. Traverse, M. D. Strand, J. J. Borges, D. B. Knoester, A. Reba, A. G. Meyer, Identifying structural variation in haploid microbial genomes from short-read resequencing data using breseq. BMC Genomics 15, 1039 (2014).

52. D. E. Deatherage, J. E. Barrick, “Identification of Mutations in Laboratory-Evolved Microbes from Next-Generation Sequencing Data Using breseq” in Engineering and Analyzing Multicellular Systems: Methods and Protocols, L. Sun, W. Shou, Eds. (Springer, New York, NY, 2014; 10.1007/978-1-4939-0554-6_12), pp. 165–188.

53. D. E. Deatherage, C. C. Traverse, L. N. Wolf, J. E. Barrick, Detecting rare structural variation in evolving microbial populations from new sequence junctions using breseq. Front. Genet. 5 (2015).

54. J. Koff, CYstic Fibrosis bacterioPHage Study at Yale (CYPHY): A Single-site, Randomized, Double-blind, Placebo-controlled Study of Bacteriophage Therapy YPT-01 for Pseudomonas Aeruginosa Infections in Adults With Cystic Fibrosis (NCT04684641) (2021).

55. A. M. Luján, S. Paterson, E. Hesse, L. M. Sommer, R. L. Marvig, M. D. Sharma, E. O. Alseth, O. Ciofu, A. M. Smania, S. Molin, H. K. Johansen, A. Buckling, Polymicrobial infections can select against Pseudomonas aeruginosa mutators because of quorum-sensing trade-offs. Nat Ecol Evol 6, 979–988 (2022).

56. R. E. Cecil, E. Ornelas, A. Phan, N. O. Medina-Chavez, M. Travisano, D. R. Yoder-Himes, Long-term culturing of Pseudomonas aeruginosa in static, minimal nutrient medium results in increased pyocyanin production, reduced biofilm production, and loss of motility. Applied and Environmental Microbiology 91, e00975–25 (2025).

57. E. Butaitė, J. Kramer, S. Wyder, R. Kümmerli, Environmental determinants of pyoverdine production, exploitation and competition in natural Pseudomonas communities. Environmental Microbiology 20, 3629–3642 (2018).

58. M. Klausen, A. Heydorn, P. Ragas, L. Lambertsen, A. Aaes-Jørgensen, S. Molin, T. Tolker-Nielsen, Biofilm formation by Pseudomonas aeruginosa wild type, flagella and type IV pili mutants. Molecular Microbiology 48, 1511–1524 (2003).

59. L. Chen, X. Zhao, S. Wongso, Z. Lin, S. Wang, Trade-offs between receptor modification and fitness drive host-bacteriophage co-evolution leading to phage extinction or co-existence. ISME J 18, wrae214 (2024).

60. F. Pourhasanzade, S. Iyer, J. Tjendra, L. Landor, S. Våge, Individual-based model highlights the importance of trade-offs for virus-host population dynamics and long-term co-existence. PLOS Computational Biology 18, e1010228 (2022).

61. J. Gurney, S. P. Brown, O. Kaltz, M. E. Hochberg, Steering Phages to Combat Bacterial Pathogens. Trends in Microbiology 28, 85–94 (2020).

62. A. Oromí-Bosch, J. D. Antani, P. E. Turner, Developing Phage Therapy That Overcomes the Evolution of Bacterial Resistance. Annual Review of Virology 10, 503–524 (2023).

63. A. Persat, Y. F. Inclan, J. N. Engel, H. A. Stone, Z. Gitai, Type IV pili mechanochemically regulate virulence factors in Pseudomonas aeruginosa. Proceedings of the National Academy of Sciences 112, 7563–7568 (2015).

64. Y. J. Huang, J. J. LiPuma, The Microbiome in Cystic Fibrosis. Clin Chest Med 37, 59–67 (2016).

65. N. S. B. Houpt, R. Kassen, On the De Novo Emergence of Ecological Interactions during Evolutionary Diversification: A Conceptual Framework and Experimental Test. The American Naturalist 202, 800–817 (2023).

66. B. N. J. Watson, E. Pursey, S. Gandon, E. R. Westra, Transient eco-evolutionary dynamics early in a phage epidemic have strong and lasting impact on the long-term evolution of bacterial defences. PLOS Biology 21, e3002122 (2023).

67. P. B. Rainey, M. Travisano, Adaptive radiation in a heterogeneous environment. Nature 394, 69–72 (1998).

68. E. Peeters, H. J. Nelis, T. Coenye, Comparison of multiple methods for quantification of microbial biofilms grown in microtiter plates. Journal of Microbiological Methods 72, 157– 165 (2008).

69. D. Hyatt, G.-L. Chen, P. F. LoCascio, M. L. Land, F. W. Larimer, L. J. Hauser, Prodigal: prokaryotic gene recognition and translation initiation site identification. BMC Bioinformatics 11, 119 (2010).

70. S. R. Eddy, Accelerated Profile HMM Searches. PLOS Computational Biology 7, e1002195 (2011).

71. M. Steinegger, J. Söding, MMseqs2 enables sensitive protein sequence searching for the analysis of massive data sets. Nat Biotechnol 35, 1026–1028 (2017).

72. J. Huerta-Cepas, D. Szklarczyk, D. Heller, A. Hernández-Plaza, S. K. Forslund, H. Cook, D. R. Mende, I. Letunic, T. Rattei, L. J. Jensen, C. von Mering, P. Bork, eggNOG 5.0: a hierarchical, functionally and phylogenetically annotated orthology resource based on 5090 organisms and 2502 viruses. Nucleic Acids Res 47, D309–D314 (2019).

73. B. Buchfink, K. Reuter, H.-G. Drost, Sensitive protein alignments at tree-of-life scale using DIAMOND. Nat Methods 18, 366–368 (2021).

74. C. P. Cantalapiedra, A. Hernández-Plaza, I. Letunic, P. Bork, J. Huerta-Cepas, eggNOG-mapper v2: Functional Annotation, Orthology Assignments, and Domain Prediction at the Metagenomic Scale. Mol Biol Evol 38, 5825–5829 (2021).

75. M. Blum, A. Andreeva, L. C. Florentino, S. R. Chuguransky, T. Grego, E. Hobbs, B. L. Pinto, A. Orr, T. Paysan-Lafosse, I. Ponamareva, G. A. Salazar, N. Bordin, P. Bork, A. Bridge, L. Colwell, J. Gough, D. H. Haft, I. Letunic, F. Llinares-López, A. Marchler-Bauer, L. Meng-Papaxanthos, H. Mi, D. A. Natale, C. A. Orengo, A. P. Pandurangan, D. Piovesan, C. Rivoire, C. J. A. Sigrist, N. Thanki, F. Thibaud-Nissen, P. D. Thomas, S. C. E. Tosatto, C. H. Wu, A. Bateman, InterPro: the protein sequence classification resource in 2025. Nucleic Acids Res 53, D444–D456 (2025).

76. T. Paysan-Lafosse, A. Andreeva, M. Blum, S. R. Chuguransky, T. Grego, B. L. Pinto, G. A. Salazar, M. L. Bileschi, F. Llinares-López, L. Meng-Papaxanthos, L. J. Colwell, N. V. Grishin, R. D. Schaeffer, D. Clementel, S. C. E. Tosatto, E. Sonnhammer, V. Wood, A. Bateman, The Pfam protein families database: embracing AI/ML. Nucleic Acids Res 53, D523–D534 (2025).

77. G. Yu, L.-G. Wang, Y. Han, Q.-Y. He, clusterProfiler: an R Package for Comparing Biological Themes Among Gene Clusters. OMICS: A Journal of Integrative Biology 16, 284–287 (2012).

78. T. Wu, E. Hu, S. Xu, M. Chen, P. Guo, Z. Dai, T. Feng, L. Zhou, W. Tang, L. Zhan, X. Fu, S. Liu, X. Bo, G. Yu, clusterProfiler 4.0: A universal enrichment tool for interpreting omics data. Innovation 2 (2021).

79. S. Xu, E. Hu, Y. Cai, Z. Xie, X. Luo, L. Zhan, W. Tang, Q. Wang, B. Liu, R. Wang, W. Xie, T. Wu, L. Xie, G. Yu, Using clusterProfiler to characterize multiomics data. Nat Protoc 19, 3292– 3320 (2024).

80. G. Yu, Thirteen years of clusterProfiler. Innovation 5 (2024).

81. Y. Benjamini, Y. Hochberg, Controlling the False Discovery Rate: A Practical and Powerful Approach to Multiple Testing. Journal of the Royal Statistical Society: Series B (Methodological) 57, 289–300 (1995).

