## Supplemental Materials for "Community diversity favors coexistence between bacteria and parasitic bacteriophages"

### Supplementary Materials

#### Colony Morphology

During experimental evolution, we observed colony morphologies indicative of the emergence and maintenance of intrapopulation diversity in our experimentally evolving *Pseudomonas* populations. Pseudomonas colonies were counted by plating onto selective Pseudomonas-isolation agar (see Methods). During this plating, we observed the emergence of a novel small colony morphology (Fig S1). Plated volumes were too small to reliably estimate the exact frequency of small colony variants, but there is a trend of populations in the phage-added treatment initially evolving the small colony variant later than those in the no-phage treatment. The evolution and maintenance of within-population colony morphology diversity under static culture conditions is consistent with much past work on the topic (*67*). However, note that these colony morphologies were only apparent on Pseudomonas isolation agar and not on Bolton Broth base agar, so their relevance in the selection environment is unclear.

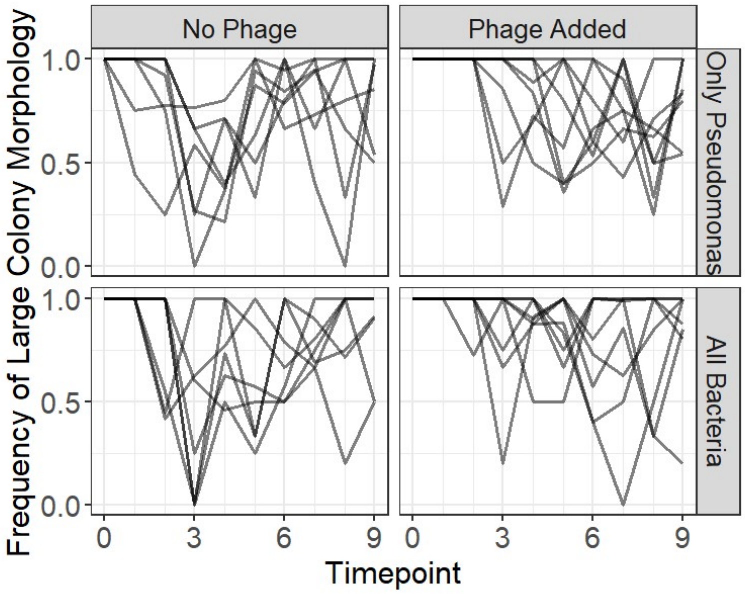


**Figure S1. *Pseudomonas* colony morphology dynamics during experimental evolution reveal extant diversity in all treatments.** During experimental evolution in Bolton Broth Base, communities were plated on Pseudomonas isolation agar to estimate *Pseudomonas* population densities. In all populations, a novel small colony morphology evolved and coexisted with the ancestral large colony morphology. Each line is an independently evolving population.

#### Preliminary Experiments

In the main text, we report the results of a single evolution experiment. However, prior to that experiment we ran two preliminary experiments under the same conditions, but with fewer replicates and passages and with the *Pseudomonas*-only and all-bacteria treatments run at different times. Although this makes rigorous direct comparison for the effect of community context impossible, the results were qualitatively similar enough to warrant inclusion. In the *Pseudomonas*-only treatment, several phage populations reached or trended towards extinction by the end of the experiment (Fig S2). In contrast, in the all-bacteria treatment, all phage populations persisted at high densities (Fig S3). Additionally, we observed a similar pattern of colony morphology diversification as Fig S1 among Pseudomonas colonies plated on Pseudomonas-isolation agar (Figs S4, S5), although a small colony morph never evolved in the phage-added treatment of the all-bacteria experiment (Fig S5).

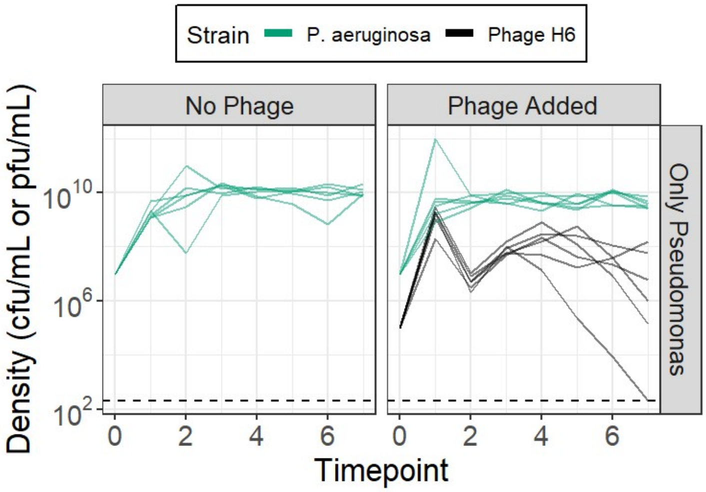


**Figure S2. Population dynamics during experimental evolution reveal that many phage populations trend towards extinction in the absence of bacterial competitors.** Bacterial and phage populations were passaged daily for 7 days with a 1:100 dilution, with population densities measured by plating. For each strain, each line is an independently evolving population.

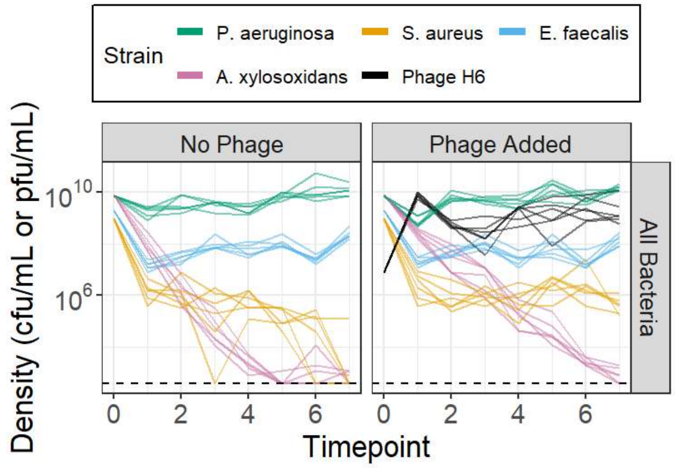


**Figure S3. Population dynamics during experimental evolution reveal that phage populations persist in the all-bacteria treatment.** Bacterial and phage populations were passaged daily for 7 days with a 1:100 dilution, with population densities measured by plating. For each strain, each line is an independently evolving population.

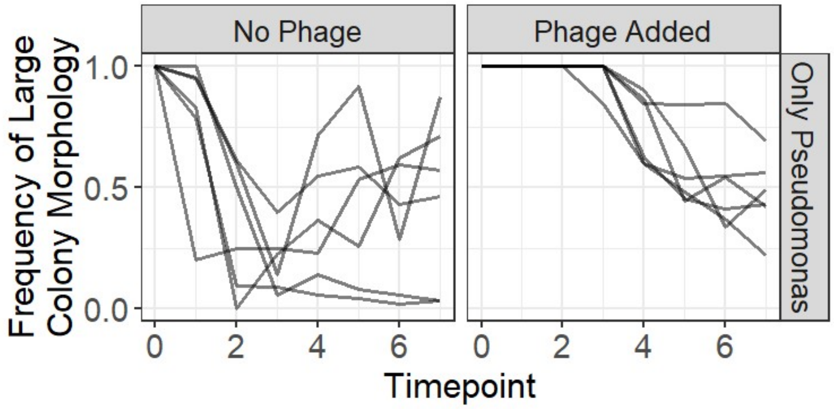


**Figure S5. *Pseudomonas* colony morphology dynamics during experimental evolution in the *Pseudomonas*-only treatment reveal extant diversity in both treatments.** During experimental evolution in Bolton Broth Base, communities were plated on Pseudomonas isolation agar to estimate *Pseudomonas* population densities. In all populations, a novel small colony morphology evolved and coexisted with the ancestral large colony morphology. Each line is an independently evolving population.

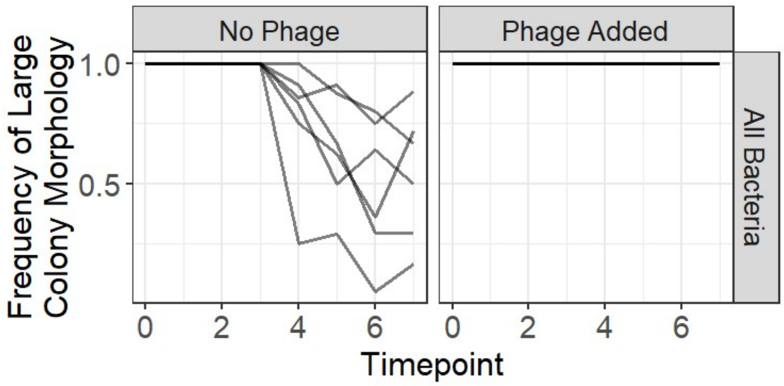


**Figure S6.** ***Pseudomonas*** **colony morphology dynamics during experimental evolution in the all-bacteria treatment reveal extant diversity in the absence of phages.** During experimental evolution in Bolton Broth Base, communities were plated on Pseudomonas isolation agar to estimate *Pseudomonas* population densities. In all populations evolved in the absence of phages, a novel small colony morphology evolved and coexisted with the ancestral large colony morphology. Each line is an independently evolving population.

#### Resistance/Infectivity Evolution

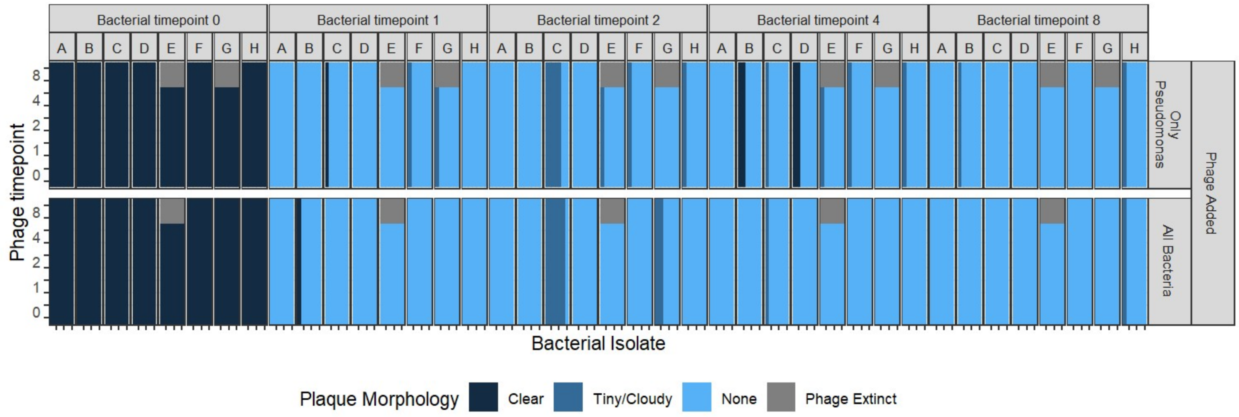


**Figure S7. Plaque morphology matches infectivity/resistance dynamics of phage H6 and *P. aeruginosa*.** Six bacterial and two phage clones were isolated from each population (A through H) at timepoints 0, 1, 2, 4, and 8. The susceptibility/resistance of all 5116 combinations was quantified, with timepoints where no phage could be isolated due to phage extinction plotted in gray.

#### Sequencing of isolates

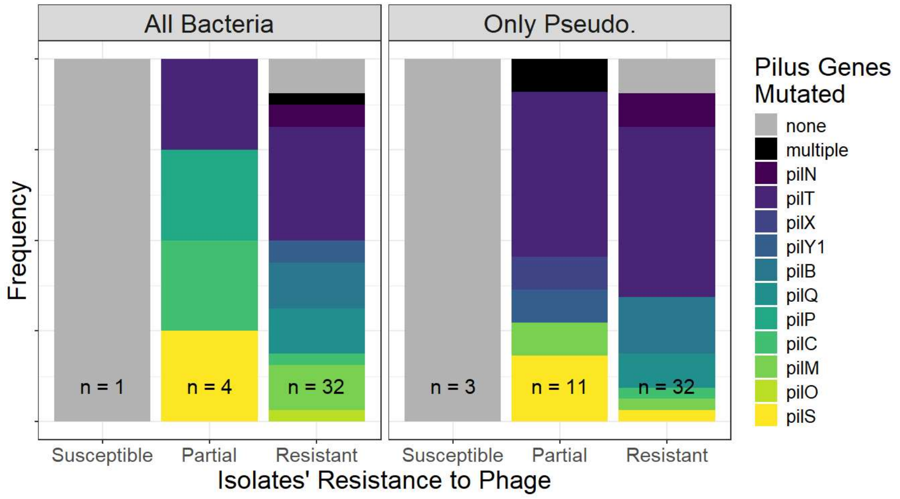


**Figure S8. Evolved phage-resistant *P. aeruginosa* clones almost always have mutations in pilus gene(s).** One evolved bacterial isolate of each resistance phenotype from each phage-added population at timepoints 1, 2, 4 and 8 was whole-genome sequenced to identify mutations. 30 of the isolates had additional mutations at other non-pilus loci (Table S1). Resistance was categorized by quantitative efficiency of plaquing and phage plaque morphology.

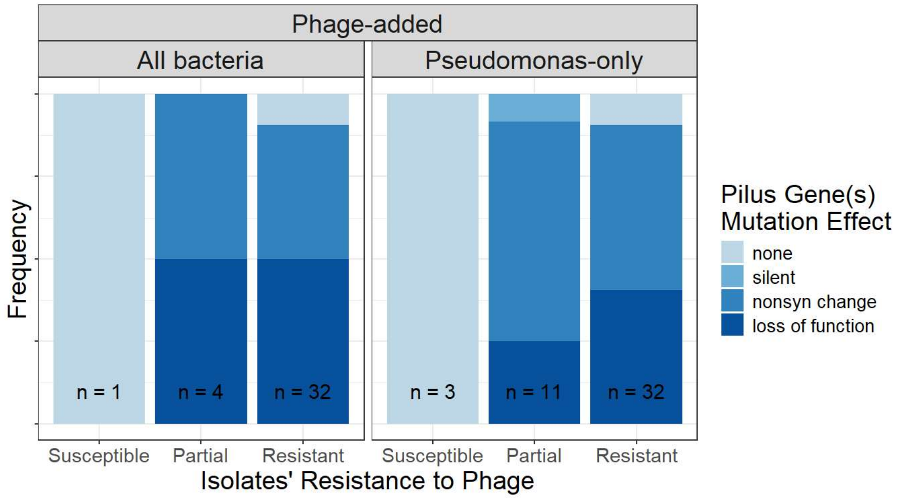


**Figure S9. Pilus mutations in evolved phage-resistant *P. aeruginosa* clones are almost always nonsynonymous or loss-of-function mutations.** One evolved bacterial isolate of each resistance phenotype from each phage-added population at timepoints 1, 2, 4 and 8 was whole-genome sequenced to identify mutations. Resistance was categorized by quantitative efficiency of plaquing and phage plaque morphology. Loss of function mutations included nonsense SNPs, small frameshift indels, and a large deletion.

[Table S1 file attached]

**Table S1. Evolved *P. aeruginosa* clones frequently have mutations in pilus and biofilm-related genes.** One evolved bacterial isolate of each resistance phenotype from each phage-added population at timepoints 1, 2, 4 and 8 was whole-genome sequenced to identify mutations. Resistance was categorized by quantitative efficiency of plaquing and phage plaque morphology.

### Bacterial isolate phenotypes

#### Materials and Methods

##### Biofilm formation

To quantify biofilm formation (*68*), bacterial isolates were grown overnight shaking in 10 mL Bolton Broth at 37°C. Cultures were diluted 1:100 in Bolton Broth and 100 µL of each added to a well of a 96 well plate and incubated statically for 24 hours at 37°C. The plate was inverted and emptied and washed with water twice then 125 µL of 0.1% crystal violet was added to each well and incubated at room temperature for 15 minutes. The plate was then inverted and emptied and washed with water four times, before being blotted vigorously onto paper towels and dried overnight to remove remaining liquid. Finally, 125 µL of 30% acetic acid was added to each well and incubated at room temperature for 15 minutes. 100 µL of this solution was moved to a new 96 well plate, and the absorbance was measured at 550 nm. We quantified biofilm formation for two isolates from each population and each treatment at timepoints 1, 2, 4, and 8. In each batch there were two technical replicates of the pilA control strain, and a single technical replicate of every other strain. Batch correction was carried out by mixed-effects modeling $\log_{10}$ absorbance values by fixed effects of treatment and timepoint plus random effects of population, isolate, and batch, then subtracting the batch effect.

##### Pyocyanin and pyoverdine production

To quantify pyocyanin and pyoverdine production, bacterial isolates were grown overnight shaking in 10 mL Bolton Broth at 37°C. 100 µL of each was added to 10 mL of Bolton Broth in a 25 mL flask and incubated statically at 37°C for 24 hours. Flasks were shaken for 10 minutes at 37°C, then the contents were transferred into a Falcon tube. Falcon tubes were spun at maximum speed for 20 minutes, then supernatant was removed and passed through a 0.2 µm filter. 200 uL of each was added to a 96 well plate and the absorbance at 405 nm (pyoverdine) and 695 nm (pyocyanin) was measured. Batch correction was carried out by mixed-effects modeling absorbance at each wavelength by fixed effects of treatment and timepoint plus random effects of population, isolate, and batch, then subtracting the batch effect.

##### Multivariate analyses

To analyze bacterial evolution of traits in multivariate space, we carried out principal component analysis (Fig S14) on batch-corrected values of biofilm formation and pyocyanin and pyoverdine production, as well as lag time, maximum growth rate, maximum density, and area under the curve. We excluded all incomplete cases (where a bacterial isolate was missing data for any of the seven traits), and excluded all data from population 9 since both no-phage treatments became contaminated.

#### Results

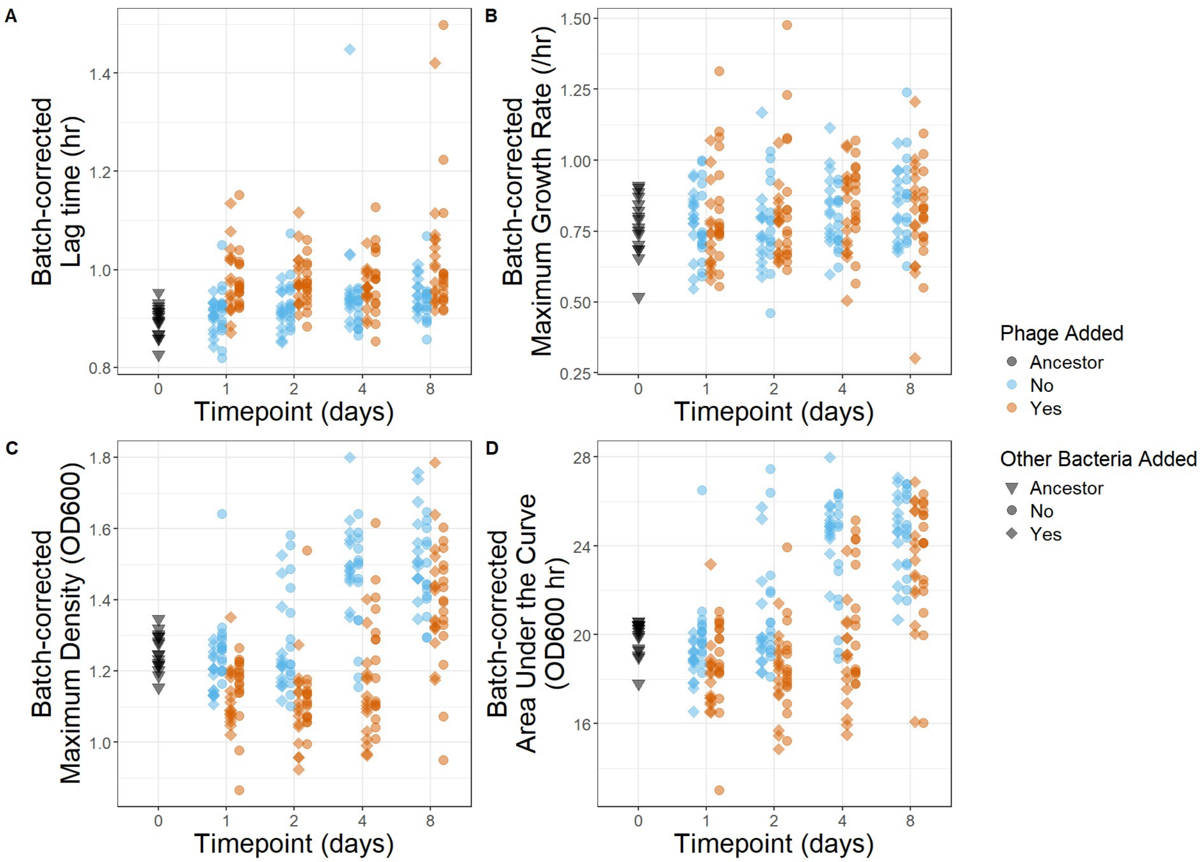


**Figure S10. *P. aeruginosa* in phage-added treatments show phenotypic costs of resistance.** For each of two evolved clones from each population and timepoint, we used growth curves to quantify lag time (A), maximum growth rate (B), maximum density (C), and area under the curve (D). Note that S10A and S10C reproduce Fig 3C and 3D, and are included here for ease of comparison.

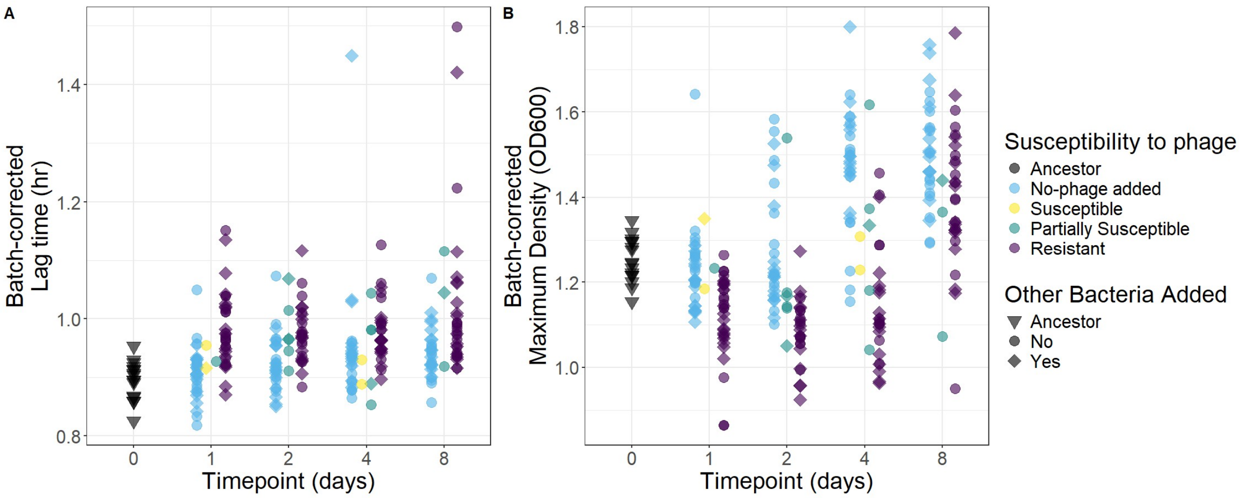


**Figure S11. *P. aeruginosa* show phenotypic costs of resistance.** For each of two evolved clones from each population and timepoint, we used growth curves to quantify lag time (A) and maximum density (B). Resistance was categorized by quantitative efficiency of plaquing and phage plaque morphology.

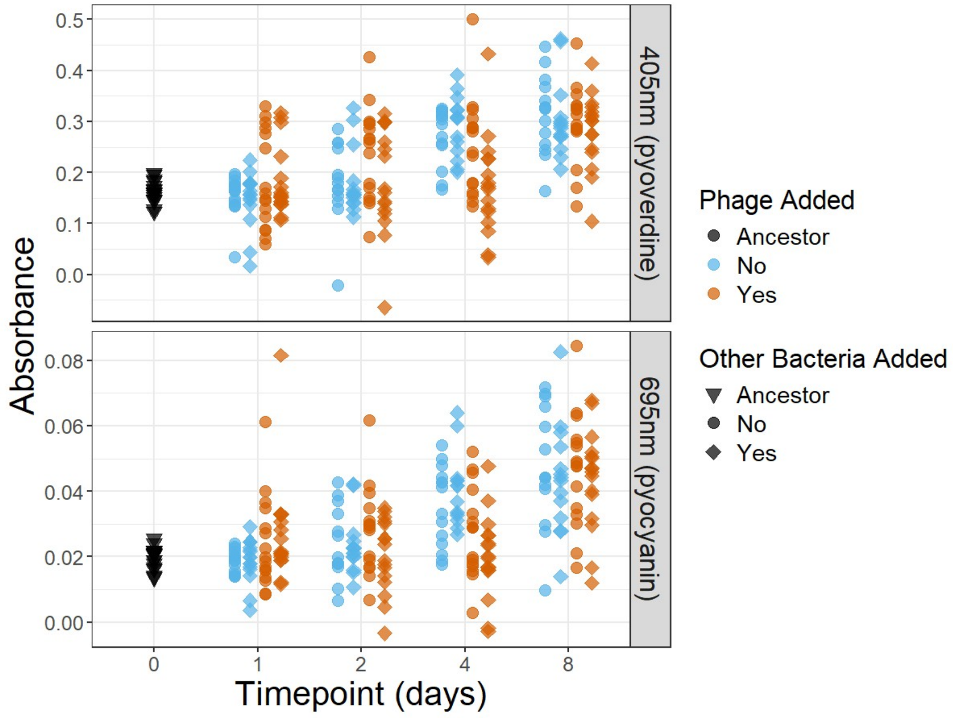


**Figure S12. *P. aeruginosa* show overall increases in pyoverdine and pyocyanin production, regardless of treatment.** For each of two evolved clones from each population and timepoint, we used spectrophotometry to quantify the excretion of two Pseudomonas-excreted pigments: pyoverdine (absorbance measured at 405 nm) and pyocyanin (absorbance measured at 695 nm).

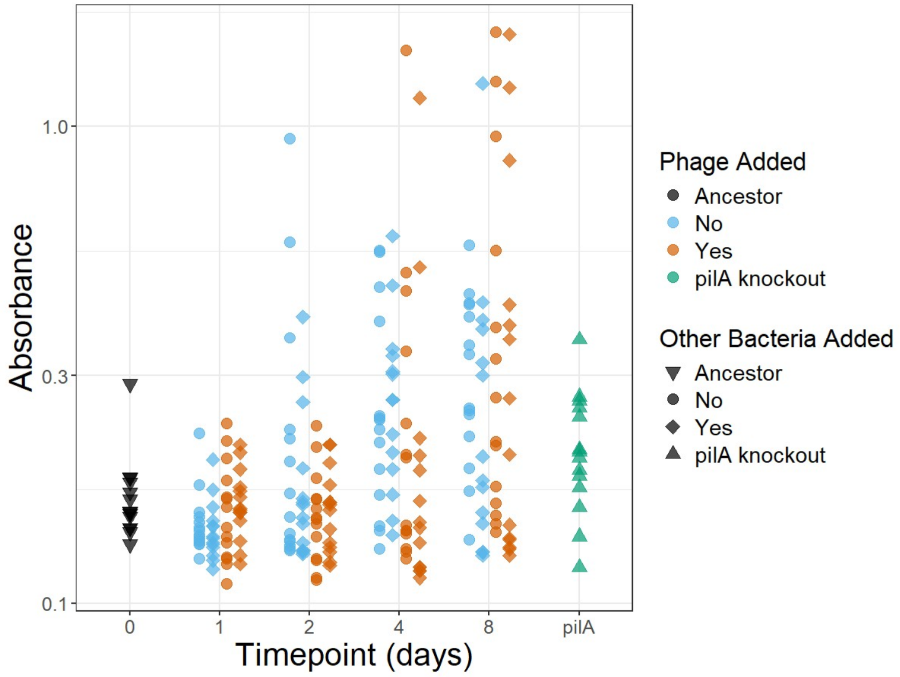


**Figure S13. *P. aeruginosa* show overall increases in biofilm formation, regardless of treatment.** For each of two evolved clones from each population and timepoint, we quantified biofilm formation by crystal violet staining (see Methods). Note the log-scale y axis for visualization of highly dispersed absorbance values.

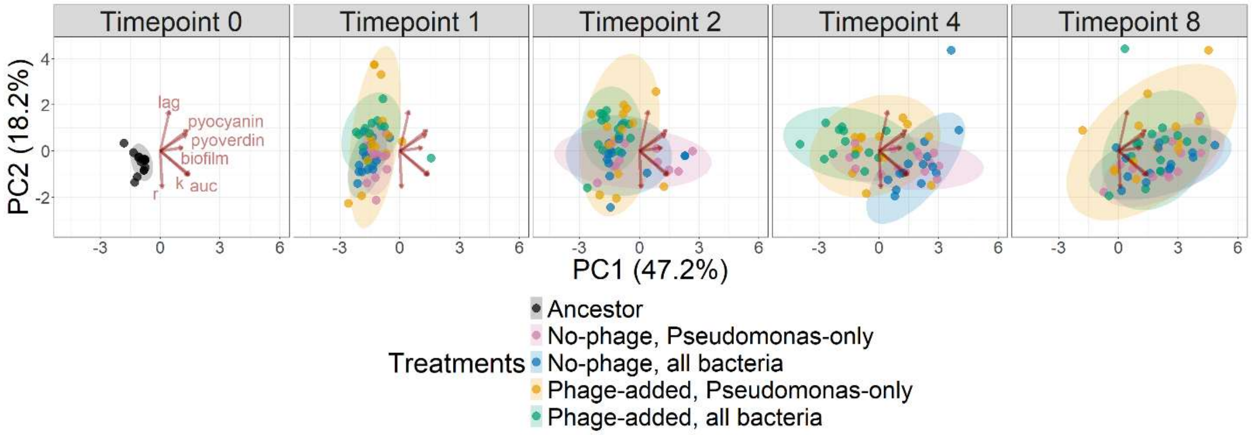


**Figure S14.** ***P. aeruginosa*** **in phage-added treatments show higher diversity and phenotypic costs of resistance.** For each of two evolved clones from each population and timepoint, we quantified biofilm formation and pyocyanin and pyoverdine production, as well as lag time, maximum growth rate ('r'), maximum density ('k'), and area under the curve ('auc'). Shown is a distance biplot of the first two components from a principal component analysis of the seven traits. All treatments show a shared pattern of evolving increased biofilm formation, pyocyanin and pyoverdine production, maximum density, and area under the curve over time relative to the ancestral phenotype. Bacteria in phage-added treatments evolved higher lag times and lower maximum growth rates, maximum densities, and area under the curves than in no-phage treatments. Bacteria in phage-added treatments also diversified more strongly and more rapidly than bacteria in no-phage treatments.

#### Sequencing of populations

We carried out metagenomic sequencing of our whole communities. Coverage (Fig S15) was calculated with the SAMtools v1.21-GCC-12.2.0 coverage command on breseq-output bam files. We achieved high coverage of *P. aeruginosa*, always greater than 100x, but variable and lower coverage of the other members of the community (Fig S15). Because of this, we focused on evolution in *P. aeruginosa*, especially on mutations at frequencies above 0.1.

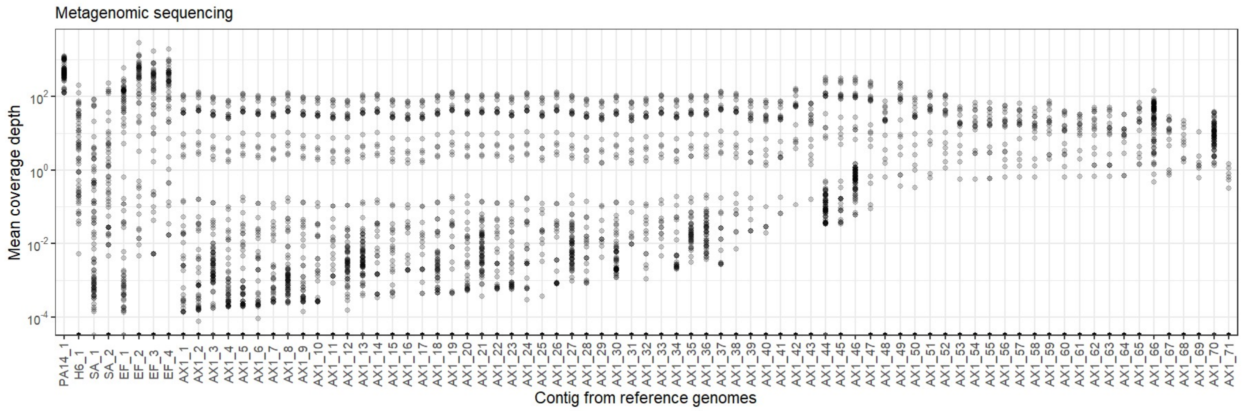


**Figure S15. Metagenomic sequencing yielded high coverage of *P. aeruginosa* and variable but lower coverage of other species.** Whole communities in no-phage treatments were sequenced at timepoints 0 and 8, while communities in phage-added treatments were sequenced at timepoints 0, 1, 2, 4, and 8. Reads were aligned to each reference genome during mutation calling by breseq.

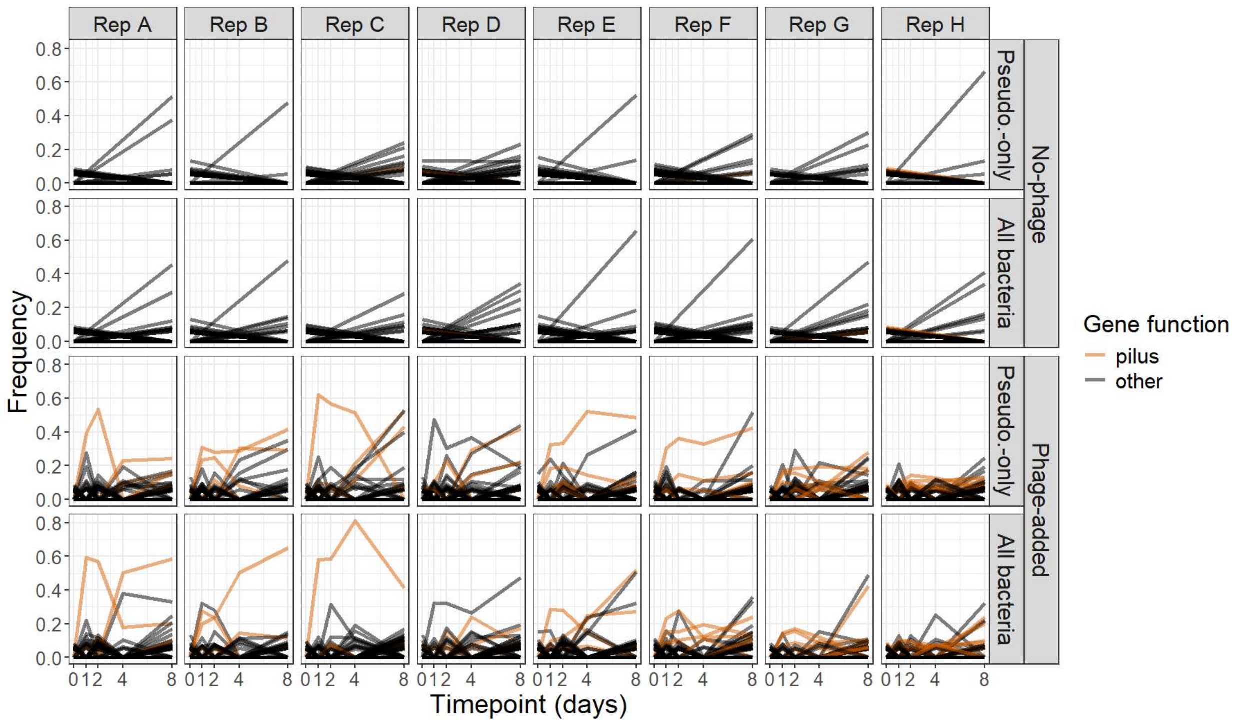


**Figure S16. Molecular evolution in populations in phage-added treatment is dominated by mutations in diverse pilus-related genes.** Whole communities in no-phage treatments were sequenced at timepoints 0 and 8, while communities in phage-added treatments were sequenced at timepoints 0, 1, 2, 4, and 8. Each line is a unique mutation observed over time. Mutations are colored by whether they affect a pilus-related gene. In contrast to Fig 4A, here all detected mutations are plotted.

Our metagenomic sequencing is short-read data, meaning that we cannot directly observe linkage between two mutations found in the same population at the same timepoint if they are further apart than the read length (150 bp). In order to calculate the total frequency of mutations affecting the pilus, we can make different assumptions about the linkage between distinct pilus mutations. In the main text, we report results where we assumed that distinct pilus mutations arise on separate genetic backgrounds and are competing with one another (an 'exclusion' assumption). Under this assumption, the total frequency of pilus mutations is simply the sum of the frequencies of each pilus mutation: $\sum f_i$. This assumption is consistent with our isolate data sequencing data (Fig S8), where very few isolates have multiple mutations in pilus-related genes. It is also consistent with the putative effect of the observed pilus mutations, where many mutations are loss of function mutations, and the remainder are all non-synonymous changes (Figs 4B, S9). Moreover, we find that this nearly-always produces a total mutation frequency between 0 and 1. However, for completeness here we also report results with a more conservative assumption: that mutations arise on a random genetic background independent of each other (an 'independence' assumption). Under this assumption, the total frequency of pilus mutations is $1 - \prod(1 - f_i)$.

As expected, pilus mutations are significantly more frequent in phage-added treatments, regardless of the linkage assumption (Fig S17, ANOVA for inclusion of phage treatment factor on fit of mixed effect model for total frequency at timepoint 8, with fixed-effect for each treatment and random effect for population, $p < 0.001$ under both exclusion or independence assumption). Generally however, the relationship between pilus mutation frequency and phage density or community treatment was weaker under the independence assumption than the exclusion assumption. Phage density was strongly and significantly negatively correlated with total pilus mutation frequency under the exclusion assumption (Fig S18A, linear model of $\log_{10}$ phage density by treatment × pilus mutation frequency, only-Pseudomonas coefficient = -3.3, p = 0.04; all-bacteria coefficient = -3.1, p = 0.05), but not the independence assumption (Fig S18B, only-Pseudomonas p = 0.08, all-bacteria p = 0.23). Pilus mutation frequencies were moderately but significantly higher in the Pseudomonas-only treatment under the exclusion assumption (Fig S18C, Pseudomonas-only effect of 0.12 with p = 0.027) but not the independence assumption (Fig S18D, Pseudomonas-only effect of 0.03 with p = 0.29).

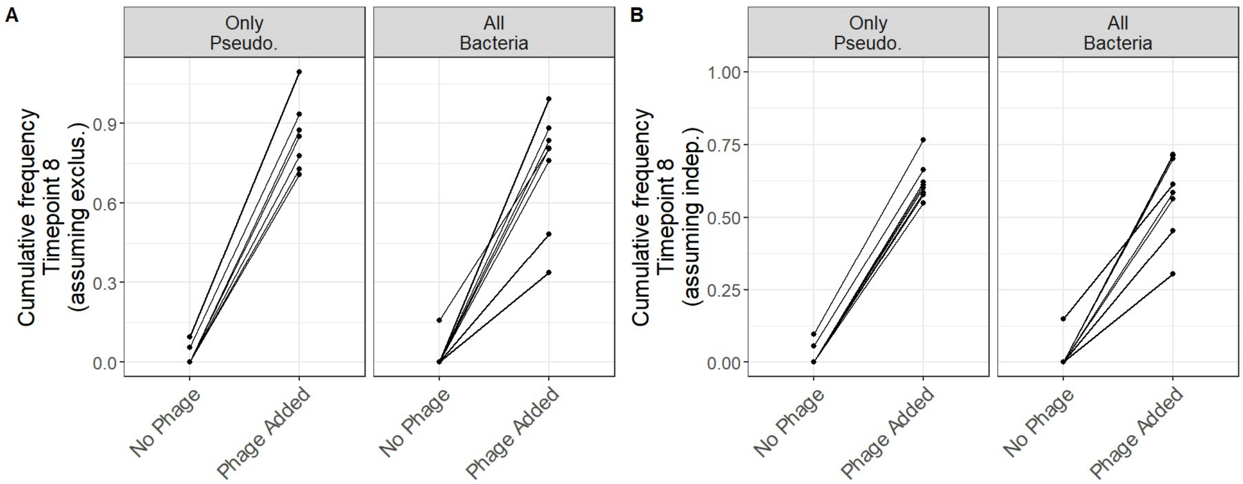


**Figure S17. Mutations in pilus-related genes are more frequent in the phage-added treatment. A.** The total frequency of all mutations in pilus-related genes, assuming that mutations arose on different genetic backgrounds and are competing with one another. **B.** The total frequency of all mutations in pilus-related genes, assuming that mutations arose independently regardless of background.

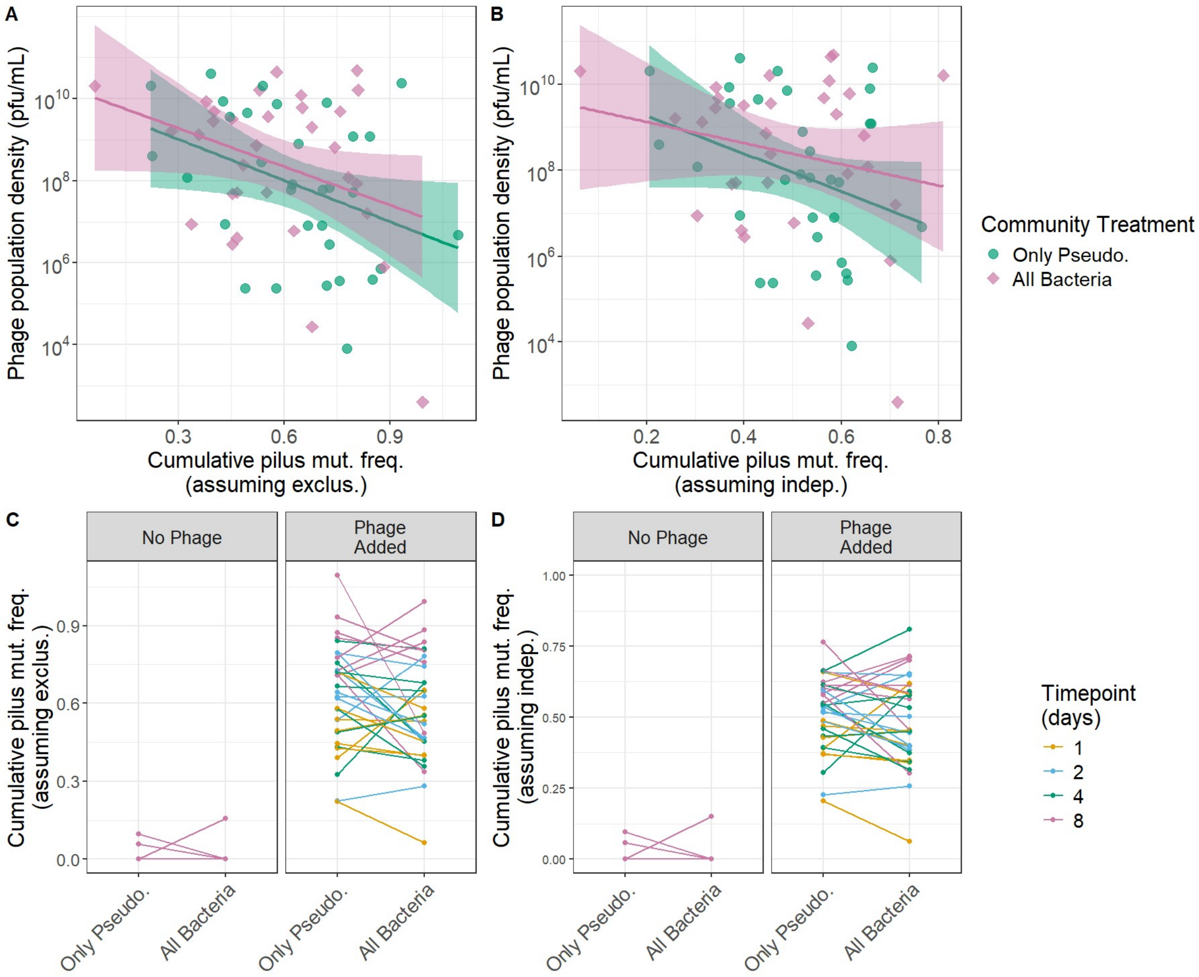


**Figure S18. Mutations in pilus-related genes predict lower phage density and are more frequent in the Pseudomonas-only treatment. A, C.** The total frequency of all mutations in pilus-related genes, assuming that mutations arose exclusively on different backgrounds. Note that S18A and S18C reproduce Fig 5A and 5B, and are included here for ease of comparison. **B, D.** The total frequency of all mutations in pilus-related genes, assuming that mutations arose independently regardless of background.

To additionally test how the phage and community treatments altered the frequency of different mutations in *P. aeruginosa*, we complemented the analyses of pilus mutation frequency reported in the main text with Gene Ontology (GO) term enrichment analysis. The *P. aeruginosa* ancestral genome was annotated with eggNOG v2.1.13 with go_evidence set to all but otherwise default settings (*69*–*74*) and InterProScan v5.77-108.0 with default settings (*75*, *76*). First, if there were multiple mutations in a single gene in a single sample, we calculated the total frequency of non-synonymous mutations in each gene conservatively by assuming independence (i.e. the distinct mutations arose on genetic backgrounds regardless of the presence of other mutations, see above for more discussion). Then, for each gene we calculated the average of the non-zero frequencies across the replicate populations in each treatment. We then calculated differences in these averaged frequencies for phage-added versus no-phage at timepoint 8, *Pseudomonas*-only versus all bacteria within the no-phage treatment at timepoint 8, and *Pseudomonas*-only versus all bacteria within the phage-added treatment at timepoints 1, 2, 4, and 8. clusterProfiler (*77*–*80*) was used to identify enriched GO terms, using a frequency difference of 0.15 as the cutoff for genes to include. Results were filtered for terms with an unadjusted p-value for enrichment ≤ 0.01 and a count of genes greater than 1 (Table S2). The analysis identified no terms enriched among mutations in the no-phage treatments, and 12 terms enriched among mutations in the phage-added treatments, including several pilus-related terms. The analysis identified no terms enriched for either community treatment within the no-phage treatment, and no terms enriched for the all-bacteria treatment within the phage-added treatment. However, within the phage-added treatment 4 terms were enriched among mutations in the Pseudomonas-only treatment relative to the all-bacteria treatment, all of which are related to the pilus. These results corroborate our findings that pilus-related mutations are more frequent in phage-added and Pseudomonas-only treatments.

**Table S2. Pilus-related GO terms are enriched in phage-added and Pseudomonas-only treatments.** Enriched Gene Ontology terms were identified using clusterProfiler from genes whose average difference in non-zero mutation frequencies was greater than 0.15. Differences in non-zero frequencies were calculated for phage-added versus no-phage among all treatments at timepoint 8, *Pseudomonas*-only versus all bacteria at timepoint 8 within the no-phage treatment, and *Pseudomonas*-only versus all bacteria at timepoints 1, 2, 4, and 8 within the phage-added treatment. Raw p-values are reported, as well as p-values corrected for multiple testing using the Benjamini & Hochberg adjustment (*81*).

| Treatment | Within treatment | GO terms | Fold Enrichment | p-value | Adjusted p-value |
|---|---|---|---|---|---|
| No-phage | All | None | - | - | - |
| Phage-added | All | pilus assembly | 71.15 | 0.00001 | 0.001 |
| | | enterobacterial common antigen biosynthetic process | 260.88 | 0.00001 | 0.001 |
| | | enterobacterial common antigen metabolic process | 260.88 | 0.00001 | 0.001 |
| | | cell projection assembly | 35.57 | 0.00007 | 0.003 |
| | | pilus organization | 31.31 | 0.00010 | 0.004 |
| | | cell projection organization | 19.57 | 0.00042 | 0.013 |
| | | peptide transport | 11.68 | 0.00191 | 0.043 |
| | | amide transport | 10.87 | 0.00235 | 0.047 |
| | | peptide secretion | 21.74 | 0.00362 | 0.066 |
| | | protein transport | 7.53 | 0.00665 | 0.108 |
| | | intracellular protein localization | 6.99 | 0.00816 | 0.108 |
| | | establishment of protein localization | 6.93 | 0.00837 | 0.108 |
| Pseud.-only | No-phage | None | - | - | - |
| All bacteria | No-phage | None | - | - | - |
| Pseud.-only | Phage-added | pilus assembly | 84.32 | 0.00023 | 0.029 |
| | | cell projection assembly | 42.16 | 0.00093 | 0.053 |
| | | pilus organization | 37.10 | 0.00121 | 0.053 |
| | | cell projection organization | 23.19 | 0.00309 | 0.063 |
| All bacteria | Phage-added | None | - | - | - |
